# Interrogating antiviral antibody responses with multiplexed, high-throughput serum assays

**DOI:** 10.64898/2026.01.03.697501

**Authors:** Amanda C. Hornick, Lucy C. Walters, Connor S. Dobson, Stephanie A. Gaglione, Michael E. Birnbaum

## Abstract

The COVID-19 pandemic underscored the importance of rapidly analyzing antibody responses against emerging viruses. Existing techniques, however, are limited in their ability to probe antibodies’ recognition of multiple native-conformation antigens simultaneously. To increase the throughput and multiplexability of antibody profiling, we developed **A**ntibody **R**eactivity **C**haracterization by **A**ntibody-**D**ependent **E**nhancement (**ARCADE**). This assay employs an antigen-agnostic Fc receptor-expressing cell line and a library of antigen-displaying, genetically barcoded lentiviruses that, when mixed with serum, infect cells and integrate their barcodes at rates reflecting the relative abundances and affinities of the antigen-specific antibodies present. Verified using sera from COVID-19-convalescent and - vaccinated donors, ARCADE delivers insights that align with and expand upon those offered by established immunoassays, highlighting, for example, how an mRNA-based vaccine elicits broader and stronger antibody responses than an adenovirus vector-based vaccine. ARCADE can comprehensively assess how infection and vaccination impact antiviral antibody repertoires over time and across patient populations.

## Introduction

The COVID-19 pandemic highlighted the significant threat that viral infections pose to human life and wellbeing. According to the World Health Organization, over 778 million people have contracted and more than 7 million people have died from COVID since it was discovered in December 2019.^1,2^ With expanding populations and rising migration levels, growing numbers of unvaccinated individuals, and changing environments introducing disease vectors to new hosts, infectious disease outbreaks from both endemic and emerging viruses are likely to increase in frequency and severity.^3^ To effectively control these outbreaks and prevent morbidity and mortality, we must urgently develop more efficient methods of measuring populations’ existing and achievable immunity against these pathogens.

Effective monitoring of humoral responses against emerging and endemic viruses requires effective antibody profiling methods. The gold standard immunoassay, the enzyme-linked immunosorbent assay (ELISA), is a sensitive, specific, and robust way to profile antibody responses, but it cannot be readily multiplexed. ELISAs are also relatively slow to implement in the context of emerging infectious diseases because they require recombinant expression of pathogen-derived antigens.^4,5^ Newer tools such as PhIP-Seq^6^ and VirScan^7^ are based upon phage display and next-generation sequencing, which enable analyses of antibody-antigen interactions in highly multiplexed formats. However, since these tools rely on linear peptide fragments produced in bacteria, they cannot identify antibodies that bind to conformationally specific epitopes or epitopes affected by eukaryotic post-translational modifications.^8–14^ Thus, such tools may not fully characterize an antibody repertoire’s pattern of antigen recognition and cannot distinguish between neutralizing and non-neutralizing antibody responses. Other multiplex immunoassays have addressed these challenges by utilizing mammalian cell lines to produce pseudovirus libraries that can measure antibodies’ ability to neutralize full-length antigens.^15–18^ Yet, such neutralization assays require prior knowledge and recombinant expression of viral entry receptors, which may not be readily attainable for emerging viruses.^15–18^

Additionally, for viruses like SARS-CoV-2, HIV, and RSV, non-neutralizing antibodies that mediate Fc effector functions also contribute to immunity.^19–23^

To overcome these limitations, we have developed a new method: **A**ntibody **R**eactivity **C**haracterization by **A**ntibody-**D**ependent **E**nhancement (**ARCADE**). This assay employs libraries of antigen-displaying, genetically barcoded lentiviruses to screen serum antibodies for their recognition of full-length, native-conformation antigens in a high-throughput, multiplexed fashion. In this assay, interactions between library lentiviruses and antigen-specific antibodies are recorded through the infection of antigen-agnostic Fc receptor-expressing cells via an antibody-dependent enhancement (ADE)-like mechanism.^24–26^ By mediating the cellular entry of antibody-lentivirus complexes, Fc receptors enable the detection of antigen-specific antibodies without the need for separate expression, or even identification, of viral entry receptors. Once the antibody-lentivirus complexes enter cells, the lentiviruses can integrate their barcodes into the cells’ genomes, enabling next-generation sequencing-based quantification of the relative strength and breadth of antigen recognition by serum antibodies.

Here, we describe ARCADE and exhibit its utility in the context of COVID-19. Using a lentiviral library displaying SARS-CoV-2 spike variants and other viral proteins, we survey sera from pre-pandemic and COVID-19-convalescent donors, demonstrating that our approach can robustly distinguish between seronegative and seropositive samples and record profiles of variant recognition that are consistent with known patterns of immune escape. We subsequently apply ARCADE to profile sera from COVID-19-vaccinated donors, illustrating how ARCADE can identify unique features of antibody responses elicited by different vaccines. We also verify our readouts against those from established immunoassays and employ the same lentiviral library in a complementary multiplexed neutralization assay to determine whether serum samples’ antibodies can both bind and neutralize library members.

With viral infectious disease outbreaks becoming more frequent,^3,27–30^ there is a pressing need for technologies that can screen antibody repertoires more efficiently. Such technologies will be critical for the rapid assessment of populations’ existing antiviral immunity and the efficacy of new vaccines and therapeutics. ARCADE’s high-throughput, multiplexable, and modular nature makes it well-suited to address this need and additionally profile antibody responses across a range of diseases, including cancer, autoimmunity, and allergy.

## Results

### Hybrid pseudotyped lentiviruses mixed with serum transduce Fc receptor-expressing cells in an antigen-specific and dilution-dependent manner

Our lab previously demonstrated that hybrid pseudotyped lentiviruses, co-displaying a targeting protein and an engineered viral fusogen, VSVGmut, can be used to selectively transduce cognate receptor-expressing cells (**Figure 1A**).^31^ VSVGmut is a modified version of VSVG, which retains the native fusogen’s robust ability to facilitate membrane fusion but ablates its ability to bind to its entry receptor, LDLR.^31^ To apply this pseudotyping strategy in the context of COVID-19, we previously engineered lentiviruses to co-display SARS-CoV-2 prefusion-stabilized (S2P)^32^ spike protein variants and VSVGmut. We showed that these lentiviruses could selectively infect a B cell line expressing the SARS-CoV-1 and SARS-CoV-2 spike-specific CR3022 antibody as a membrane-bound B cell receptor.^31^ Since CR3022-expressing cells underwent efficient antigen-specific transduction by the hybrid pseudotyped lentiviruses, we hypothesized that these lentiviruses could be similarly applied to study the antigen recognition profiles of soluble antibodies in serum.

**Figure 1.**
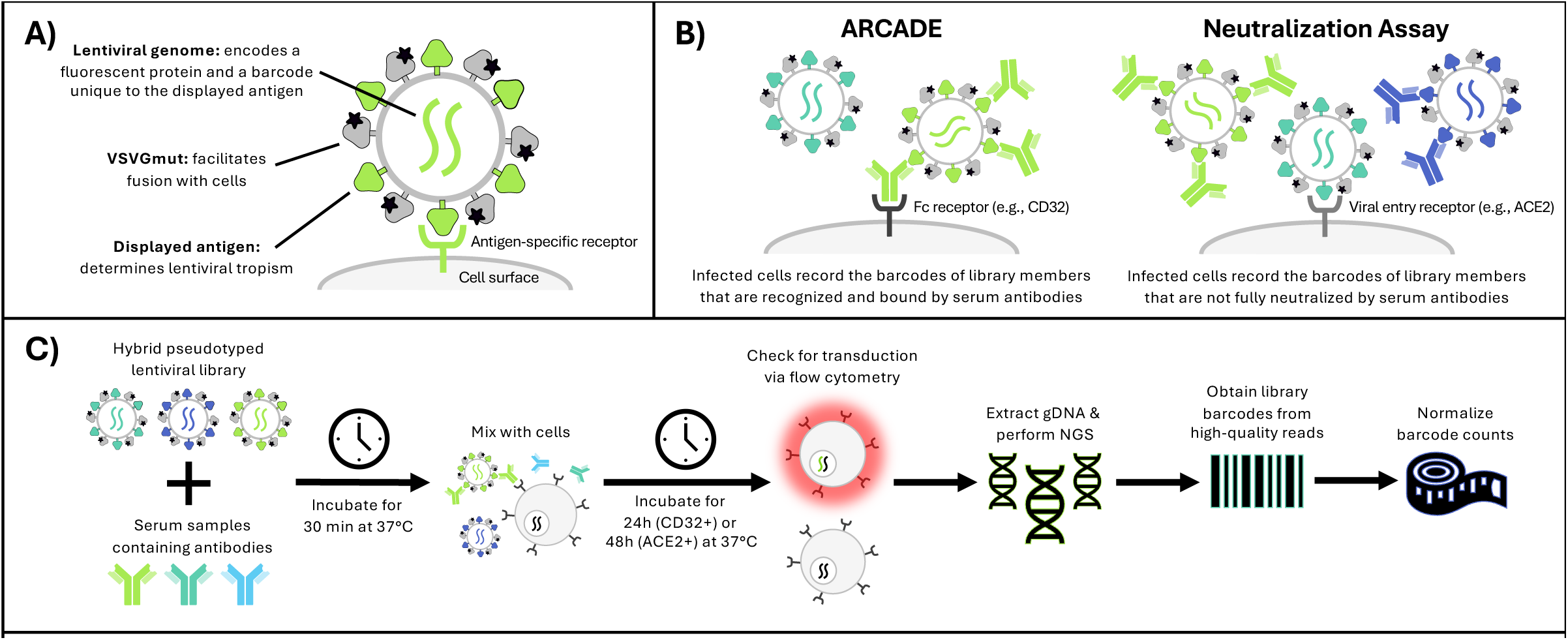
Overview of our lentivirus display-based assays. **A)** Established system for hybrid pseudotyping of lentiviruses using the VSVGmut fusogen and antigens of interest. **B)** Comparison of our high-throughput, multiplexed serum assays. ARCADE (Antibody Reactivity Characterization by Antibody-Dependent Enhancement) utilizes an Fc receptor-expressing cell line to record which lentiviral library members are bound by serum antibodies. The neutralization assay utilizes an antigen-specific viral entry receptor-expressing cell line to record which lentiviral library members are not fully neutralized by the serum antibodies. **C)** Schematic of the pipeline for our assays. The lentiviral library and serum are mixed and added to Jurkat cells expressing either CD32 for ARCADE or ACE2 for the neutralization assay. Transduction is confirmed via cells’ mCherry expression. Genomic DNA (gDNA) is extracted from assay cells, and next-generation sequencing (NGS) records genomically integrated barcodes corresponding to library members. Normalizations, described herein, are performed to allow for more robust comparisons across library members and samples.

To this aim, we engineered cells to express Fc receptors (FcRs) so that antibodies, rather than B cell receptors, could mediate the recognition and subsequent internalization of library lentiviruses. While antibody variable regions mediate antigen interactions, antibody constant regions enable downstream effector functions via FcR engagement. Similarly, FcRs facilitate antibody-dependent enhancement (ADE), internalizing antibody-virus complexes.^24–26^ Therefore, we hypothesized that FcRs could mediate the cellular internalization of lentiviruses displaying antigens recognized by serum antibodies in our assay via a mechanism similar to ADE. Thus, we termed our approach **ARCADE** (**A**ntibody **R**eactivity **C**haracterization by **A**ntibody-**D**ependent **E**nhancement) (**Figure 1B and 1C**).

We initially demonstrated the principle of ARCADE by incubating lentivirus pseudotyped with VSVGmut and wildtype (Wuhan Strain) SARS-CoV-2 S2P spike protein with a range of dilutions of sera derived from COVID-19-convalescent and pre-pandemic donors. These lentivirus-serum mixtures were added to K562 cells, which endogenously express FcγRII (CD32),^33^ and transduction rates were assessed after 24 hours via flow cytometry. While pre-pandemic donor (PPD) sera resulted in negligible transduction across all dilutions, convalescent donor (CD) sera enabled transduction in a serum dilution-dependent manner (**Figure 2A**). In the 1:40 to 1:80 dilution range, CD sera transduction rates peaked and were two-to-ten-fold higher than those for PPD sera. For CD sera dilutions below 1:40 and above 1:80, transduction rates fell substantially, and CD and PPD sera transduction rates converged at the 1:1280 dilution. Although CD32 preferentially engages aggregated, antigen-bound IgG, it is still capable of binding monomeric IgG, albeit with lower affinity.^34,35^ Therefore, we hypothesize that the hook effect observed at the lower sera dilutions results from lentivirus-bound anti-SARS-CoV-2 antibodies and unbound antibodies competing for available Fc receptors, limiting transduction.

**Figure 2.**
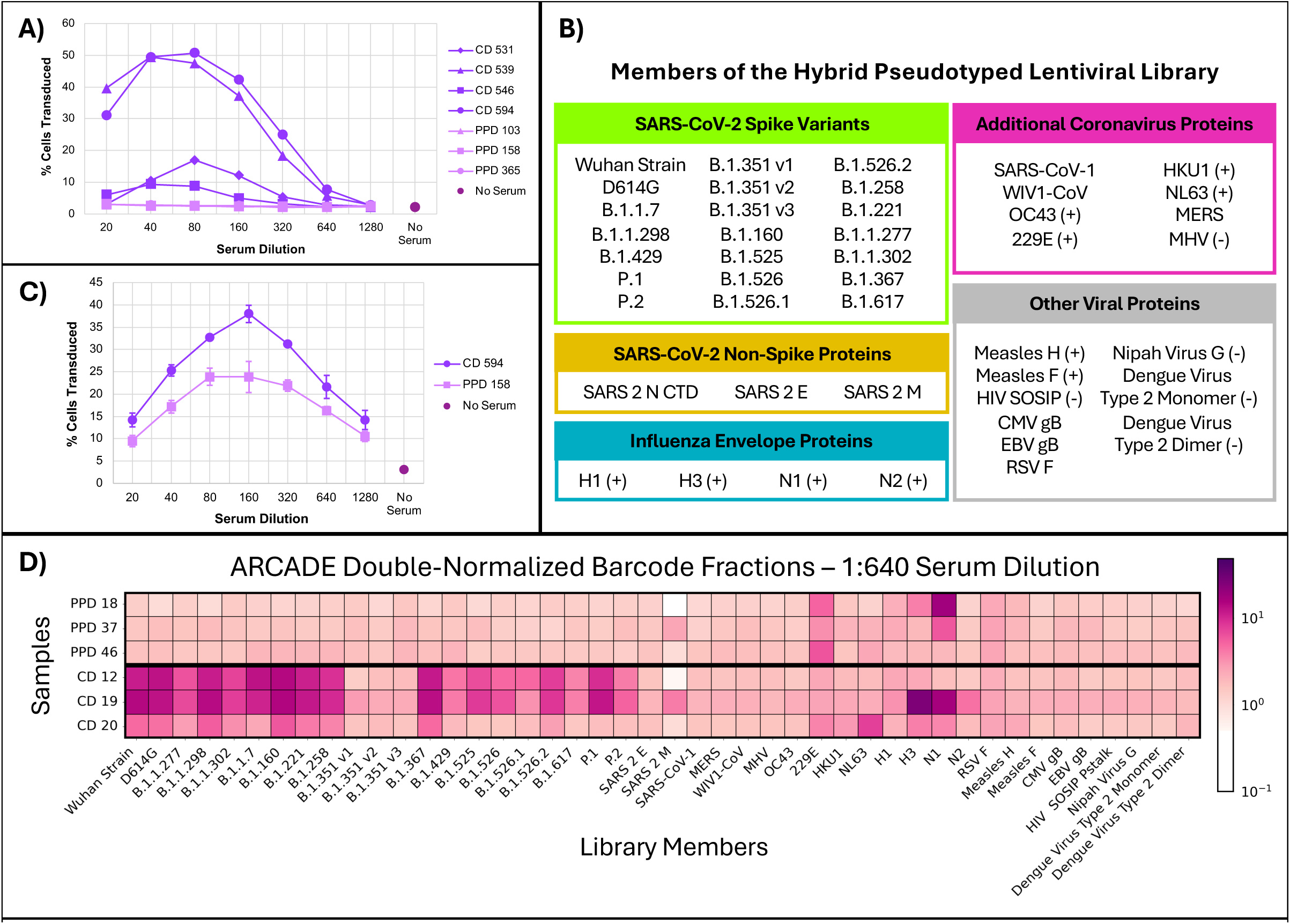
ARCADE differentiates between the viral protein recognition patterns of pre-pandemic donor (PPD) and COVID-19-convalescent donor (CD) sera. **A)** Transduction of K562 cells by a hybrid pseudotyped lentivirus in the presence of PPD or CD sera across a range of serum dilutions. Lentiviruses were pseudotyped with VSVGmut and wildtype (Wuhan Strain) SARS-CoV-2 prefusion-stabilized (S2P) spike. K562 cells express the Fc receptor FcγRII, also known as CD32. Cells were mixed with the lentivirus in the absence of serum as a control (No Serum). Averages are represented (n = 2; n = 14 for No Serum). **B)** Composition of the hybrid pseudotyped lentiviral library used for the assays performed in this work. (+) and (-) represent positive and negative controls for which most of the U.S. population is seropositive or seronegative, respectively. **C)** Transduction of CD32-expressing Jurkat cells by the hybrid pseudotyped lentiviral library in the presence of PPD or CD sera across a range of serum dilutions. Cells were mixed with the library in the absence of serum as a control (No Serum). Averages with standard deviations are represented (n = 3 for PPD and CD; n = 5 for No Serum). **D)** Double-normalized barcode fractions from ARCADE performed using the lentiviral library with PPD or CD sera diluted 1:640. Average (n = 3) double-normalized barcode fractions are represented.

We next demonstrated that ARCADE could be conducted in a multiplexed format. Libraries of different sizes were simulated by combining lentiviruses expressing either on-target (SARS-CoV-2) or off-target (HIV) antigens in different proportions and incubating the mixtures with CD and PPD sera and K562 cells. Signal-to-noise ratios, defined as the ratio of CD on-target enrichment to PPD on-target enrichment, were greatest when the proportion of on-target lentivirus used was the largest (10%) and the serum was more dilute (1:320 or 1:1280) (**Supplementary Figure 1A**). Notably, the 1% on-target lentivirus condition led to acceptable signal-to-noise ratios, suggesting that we could detect antibodies against at least 100 antigens simultaneously.

To improve ARCADE’s signal-to-noise ratio, we tested several Fc receptor-expressing cell lines for their ability to permit on-target vs off-target transduction in the presence and absence of serum. Since CD32-expressing K562 cells showed higher levels of transduction in the absence of serum than Jurkat cells (**Supplementary Figure 1B**), we generated CD32- and CD64-expressing Jurkat cell lines for use in ARCADE. We incubated these cell lines with PPD or CD serum and on-target or off-target lentiviruses and found that CD32-expressing Jurkat cells maintained on-target transduction in the presence of CD serum while exhibiting markedly reduced on-target transduction in the presence of PPD serum (**Supplementary Figure 1C**). We therefore used CD32-expressing Jurkat cells for ARCADE in place of K562 cells.

### ARCADE quantitatively profiles antiviral antibody repertoires via next-generation sequencing

We created a forty-five-member lentiviral library displaying a varied array of viral proteins (**Figure 2B** and **Supplementary Table 1**). The library included SARS-CoV-2 spike variants from the beginning of the COVID-19 pandemic through Winter 2021 as well as non-spike SARS-CoV-2 proteins.^36^ We also included proteins derived from commonly encountered viruses, such as those from influenza and measles,^37^ to serve as positive controls and proteins derived from viruses less commonly encountered by our United States-based pre-pandemic donor cohort, such as those from dengue virus and HIV, to serve as negative controls.^38,39^ The lentiviral library was assembled using our previously described LeAPS protocol^31^ to ensure that each lentiviral particle co-displays VSVGmut and a single viral antigen from the library on its surface while encoding a fluorophore (mCherry) and a unique antigen-linked barcode in its genome. As a result, lentiviral infection of assay cells and subsequent lentiviral genome integration enable both flow cytometry-based measurements of transduction and next-generation sequencing (NGS)-based readouts of relative library member infection rates.

We first validated that the forty-five-member lentiviral library also transduces CD32-expressing Jurkat cells in a serum dilution-dependent manner using flow cytometry (**Figure 2C**). As before, a hook effect was observed, with maximal transduction occurring at a dilution of 1:160 and lower levels of transduction occurring at both higher and lower dilutions. Unlike the single SARS-CoV-2 spike-displaying lentivirus, which only substantially transduced cells when mixed with CD serum, the lentiviral library showed increased transduction across dilutions of both CD and PPD serum, consistent with the library’s inclusion of positive control viral proteins. Increased transduction by the library in the presence of CD serum relative to PPD serum likely indicates the presence of SARS-CoV-2 reactive antibodies in the CD serum. Based on these data, we decided to use dilutions from the middle to the higher end of the range to maximize the signal-to-noise ratio of our assay while minimizing serum use.

To enable NGS-based readouts for ARCADE, we devised a normalization strategy that accounts for differences in serum samples’ absolute antibody levels. This strategy was key to NGS data interpretation, as without normalization, only the relative proportions of signal for each library member, but not library members’ total signal levels, could be compared between donors. To overcome this limitation and provide absolute quantifiers of signal in our assay, we added a series of three uniquely barcoded, VSVG-pseudotyped “ladder viruses” to our lentiviral library. These viruses — akin to ELISA standards — enable normalization of each serum sample’s library member barcode counts, as the viruses transduce assay cells via LDLR^31^ independently of the serum antibodies present. Therefore, samples with higher library member barcode counts relative to ladder virus barcode counts generally indicate samples with higher total antibody levels, allowing absolute antibody levels for each library member to be quantified and compared across samples (**Supplementary Figure 2A**). We utilized the ladder virus system in a two-step NGS normalization approach, enabling standardization across samples while also accounting for different rates of serum-independent background transduction across library members (**Supplementary Figure 2B**). Barcode counts for each library member were first divided by the barcode count for a given ladder virus within the same sample. Each sample’s ladder-normalized barcode fraction for each library member was then divided by the corresponding library member’s ladder-normalized barcode fraction from a measurement obtained when the assay was performed in the absence of serum.

Using this double normalization strategy, we found that CD sera consistently showed higher signal for SARS-CoV-2 variants than PPD sera (**Figure 2D**), demonstrating that ARCADE can readily distinguish between seronegative and seropositive sera. Barcode fractions were especially elevated for D614G, B.1.1.298, and B.1.160, variants which became prevalent earlier in the pandemic.^40–43^ We also observed that SARS-CoV-2 spike variants known commonly to escape humoral immunity established early in the COVID-19 pandemic (e.g., B.1.351) recorded much smaller barcode fractions than other variants in the presence of CD sera, suggesting that antibodies capable of binding such variants were largely absent from the sera.^40,43–45^ In these ways, ARCADE-based readouts aligned well with established clinical findings.

For the non-SARS-CoV-2 library members, we found that CD and PPD sera led to similar barcode fractions. Negative controls recorded negligible barcode fractions for all samples, while positive controls displaying antigens from human coronavirus 229E or influenza showed elevated barcode fractions for multiple CD and PPD serum samples (**Figure 2D**). Notably, specific donors showed markedly higher barcode fractions for 229E (PPD 18 and PPD 46), NL63 (CD 20), H3 (CD 19), and N1 (CD 19 and PPD 18) compared to other donors. The results obtained from these controls demonstrate how ARCADE can generate multiplexed profiles of antigen recognition by serum antibodies, revealing variation in antibody responses across donors and mapping potential viral exposure histories.

### ARCADE can characterize vaccine-induced antibody responses

To further demonstrate the translational utility of ARCADE and assess how vaccination affects the strength and breadth of anti-SARS-CoV-2 antibody responses, we assayed sera from donors vaccinated with either the Moderna Spikevax (mRNA-based) or the JCJ Jcovden (adenovirus vector-based) COVID-19 vaccine.^46^ Longitudinal samples were obtained from each donor across three timepoints: pre-vaccine, post dose one, and post dose two (Moderna) or pre-vaccine, two weeks post-vaccine, and two months post-vaccine (JCJ) (**Figure 3A, Supplementary Figure 3,** and **Supplementary Tables 2 and 3**).

**Figure 3.**
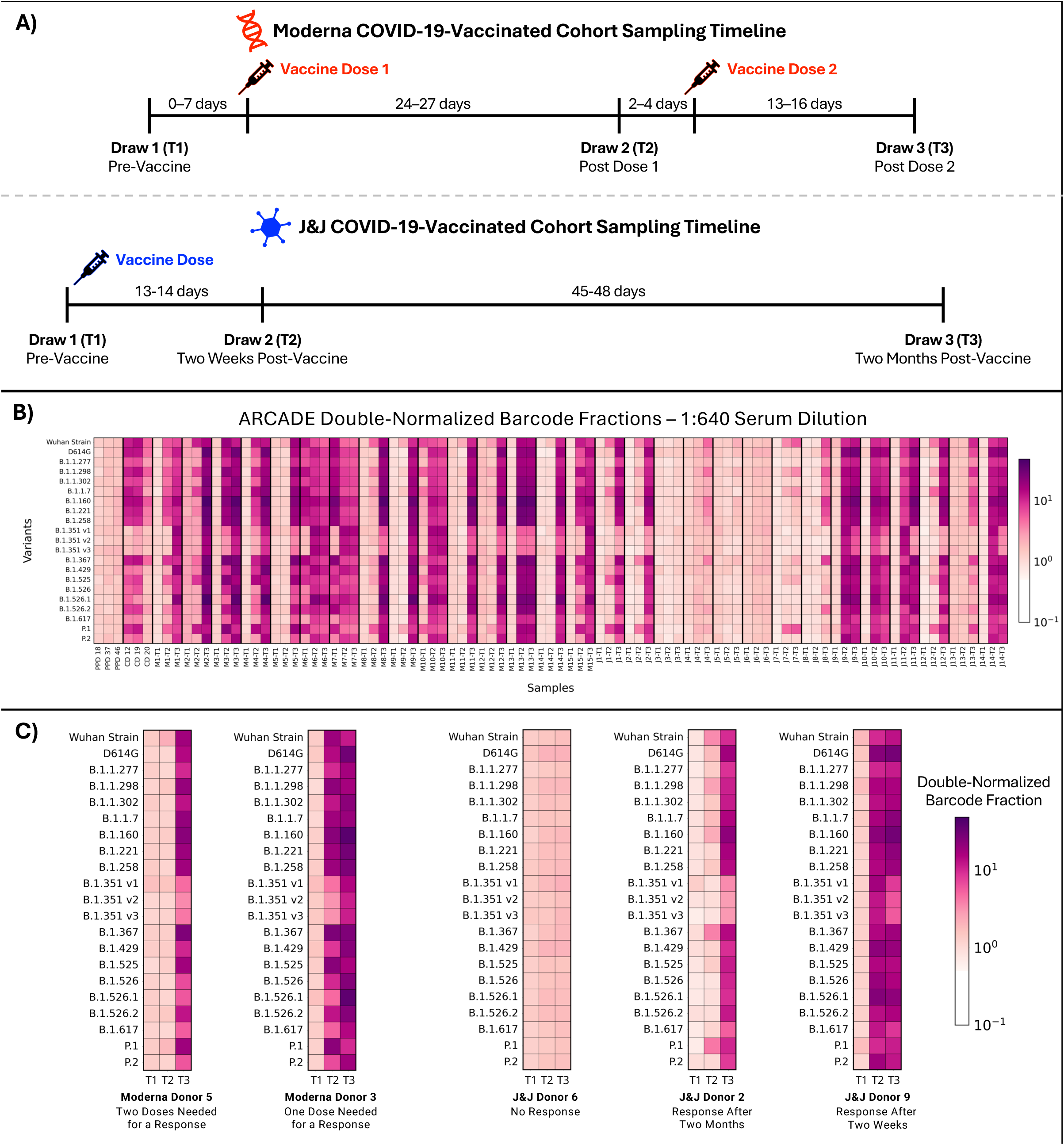
ARCADE characterizes the antiviral antibody repertoires of individuals vaccinated against COVID-19. **A)** Vaccination and blood draw timelines for the COVID-19-vaccinated cohorts. **B)** Double-normalized barcode fractions for the SARS-CoV-2 spike variants for all vaccinated and control donors. ARCADE was performed with the lentiviral library and sera — diluted 1:640 — from pre-pandemic donors (PPD), convalescent donors (CD), and vaccinated donors. Average (n = 3) double-normalized barcode fractions are shown. M = Moderna; J = JCJ. **C)** Heatmap subsections highlighting donors with different responses to the Moderna and JCJ COVID-19 vaccines. Average (n = 3) double-normalized barcode fractions are shown for serum samples diluted 1:640.

Vaccinated cohort samples were assayed via ARCADE alongside PPD sera, CD sera, and no sera as controls (**Supplementary Figure 4**). Pre-vaccine samples’ SARS-CoV-2 barcode fractions were more similar to those of the PPD sera, while post-vaccine samples’ SARS-CoV-2 barcode fractions were more similar to those of the CD sera, demonstrating that our assay can readily distinguish between donors who become seropositive and donors who remain seronegative following vaccination (**Figure 3B** and **Supplementary Figure 5**).^47–49^ Donors’ post-vaccination SARS-CoV-2 barcode fractions also highlighted how the kinetics of vaccine-induced antibody responses can differ across individuals (**Figure 3C** and **Supplementary Figure 6**). Some Moderna-vaccinated donors’ barcode fractions increased after dose one, indicating a strong response to the vaccine, while other donors’ barcode fractions increased only after dose two. Although many JCJ-vaccinated donors also exhibited increased barcode fractions post-vaccination, in contrast to the Moderna-vaccinated cohort, some JCJ-vaccinated donors did not show increased SARS-CoV-2 barcode fractions at any timepoint. These patterns align with prior reports that mRNA-based COVID-19 vaccines elicit stronger and more consistent anti-SARS-CoV-2 antibody responses than adenovirus vector-based COVID-19 vaccines.^49–53^

Our assay also enabled profiling of the unique antigen recognition and cross-reactivity patterns of vaccinated donors’ antibody repertoires. Across timepoints, most vaccinated donors exhibited smaller barcode fractions for the B.1.351 variants than for other variants (**Figure 3B**). This observation aligns with reports that the B.1.351 variants proved exceptionally resistant to neutralization — and thus binding — by serum antibodies from both convalescent and vaccinated individuals.^45,51,54^ In contrast, we observed that donors from the two vaccinated cohorts recorded the largest barcode fractions for different sets of variants post-vaccination. While Moderna-vaccinated donors’ barcode fractions were greatest for B.1.160 or B.1.526.1, JCJ-vaccinated donors’ barcode fractions were greatest for B.1.160 or earlier variants (**Supplementary Figure 6**). These results agree with those from existing clinical studies suggesting that the Moderna vaccine provided broader immunity than the JCJ vaccine, offering individuals stronger protection against variants that developed later in the pandemic.^51–53^

To better understand how donors’ antibody levels changed over time, we summed barcode fractions for the SARS-CoV-2 spike variants and control library members separately for each donor and calculated correlations between measurements from pairs of timepoints. For the SARS-CoV-2 spike variants, we observed only weak correlations (|r| < 0.38, r^2^ < 0.15) (**Supplementary Figure 7A**), but for the control library members, we observed stronger correlations (r > 0.57, r^2^ > 0.32 for Draw 2 vs Draw 1 and r > 0.69, r^2^ > 0.48 for all other comparisons) (**Supplementary Figure 7B**). These metrics suggest that donors’ pre-vaccine and mid-vaccine-series anti-SARS-CoV-2 antibody levels do not predict their post-vaccine levels and that these COVID-19 vaccines largely do not impact levels of antibodies against non-SARS-CoV-2 viral proteins.

### Established immunoassays validate ARCADE’s measurements

To validate the accuracy of the ARCADE readouts, we compared our results to both LIAISON assay antibody titers provided by the commercial serum supplier and ELISA measurements conducted in our lab. We found that there was a strong correlation (r = 0.89, p = 3.14 x 10^-21^) and positive linear association (slope = 1.91, 95% CI = [1.70-2.13]) between the ARCADE and LIASON measurements (**Figure 4**). However, this relationship between measurements did not hold for all donors. Moderna donors 6, 7, and 10 demonstrated an inverse relationship between their barcode fractions and antibody titers. These donors also had pre-vaccine antibody titers that were higher than the LIAISON assay’s cutoff for a positive result (**Supplementary Table 2** and **Supplementary Figure 3D**). Since these donors’ pre-vaccine blood draws occurred in February 2021,^36,55^ it is possible that these donors were infected with SARS-CoV-2 prior to vaccination. Post-vaccination, these donors’ SARS-CoV-2 antibody titers increased (**Supplementary Figure 3D**), but surprisingly, many of their SARS-CoV-2 variant barcode fractions decreased while their SARS-CoV-1 and WIV1-CoV barcode fractions increased substantially (**Figure 3B** and **Supplementary Figures 4 and 8**). These patterns could indicate that, in response to vaccination, these donors produced anti-SARS-CoV-2 antibodies that were highly cross-reactive with the structurally similar spikes of SARS-CoV-1 and WIV1-CoV.^56–59^ ARCADE’s highly multiplexed readouts endow it with a unique ability to identify such cross-reactive antibody responses more effectively than existing antibody-measuring techniques.

**Figure 4.**
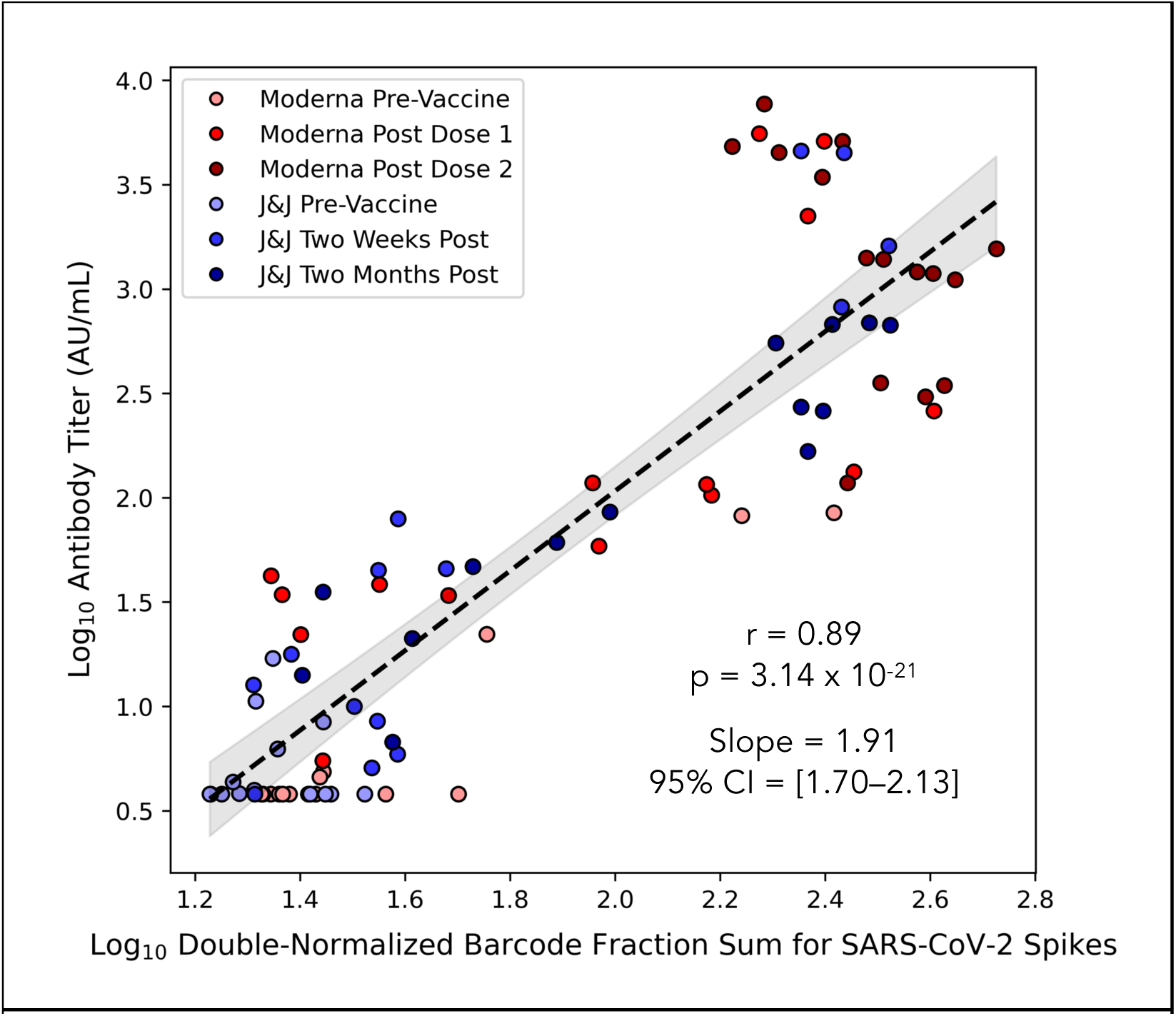
ARCADE barcode fractions and LIAISON assay antibody titers are correlated across serum samples when accounting for the inclusion of multiple measurements per donor. ARCADE was performed on vaccinated donor sera diluted 1:640. Antibody titers were provided by the serum supplier and had been measured with a LIAISON SARS-CoV-2 S1/S2 IgG assay. Double-normalized barcode fractions for all SARS-CoV-2 spike variants were summed and averaged (n = 3) for each donor at each timepoint. Both the titers and barcode fractions underwent a log_10_ transformation and were used to calculate correlation metrics across donors and timepoints to determine the overall correlation between measurements obtained with ARCADE and those obtained with the LIAISON assay. A repeated measures correlation was calculated, given that longitudinal measurements from each donor were not independent. A mixed-effects linear model was used to measure the slope of the association between the barcode fractions and antibody titers and provide the slope’s 95% confidence interval. Metrics are rounded to two places after the decimal point.

To further utilize the unique multiplexed data provided by ARCADE, we performed principal component analyses (PCA) using each donor’s library member barcode fractions as features. PCA highlighted groups of donors — in addition to Moderna-vaccinated donors 6, 7, and 10 — who shared similar patterns in both their anti-SARS-CoV-2 antibody titers and SARS-CoV-2 barcode fractions (**Supplementary Figure 9**), suggesting that our assay could stratify individuals by their immune responses to vaccination. PCA also showed, however, that donors across cohorts could have closely related library member recognition profiles, suggesting that which vaccine an individual receives is not the only factor shaping their antibody response.

When we further validated ARCADE’s measurements using anti-SARS-CoV-2 spike ELISAs for the Wuhan Strain and Alpha (B.1.1.7) variant, we generally found that serum samples with undetectable antibody titers via ELISA had ARCADE measurements under certain cutoffs (double-normalized barcode fractions of 2.10 for the Wuhan Strain and 2.11 for the Alpha variant), while samples with detectable titers had ARCADE measurements above these cutoffs at both dilutions (Supplementary Figure 10). PPD and pre-vaccine samples were among those with undetectable titers and sub-cutoff barcode fractions, while CD and post-vaccine samples had higher titers that tended to accompany larger barcode fractions. Whether the 1:160 or 1:640 dilution barcode fractions trended more closely with the ELISA titers seemed to depend on both the sample and spike variant tested.

### Hybrid pseudotyped lentiviruses can be used to perform multiplexed neutralization assays analyzed via next-generation sequencing

While ARCADE can measure antibodies that bind library-encoded antigens, it cannot determine whether those antibodies neutralize their targets. To address this limitation, we employed our lentiviral library to perform multiplexed neutralization assays. First, we engineered a Jurkat cell line to express the SARS-CoV-2 entry receptor ACE2 and tested whether the cell line could be infected by SARS-CoV-2 spike-displaying lentiviruses. We observed that ACE2+ cells were efficiently transduced by lentiviruses co-displaying VSVGmut and the Wuhan Strain SARS-CoV-2 S2P spike but not by lentiviruses displaying VSVGmut or spike alone (**Figure 5**). We also saw that ACE2- cells were not substantially transduced by any of the lentiviruses tested, confirming that we could employ ACE2+ Jurkat cells to measure rates of antigen-directed infection by hybrid pseudotyped lentiviruses.

**Figure 5.**
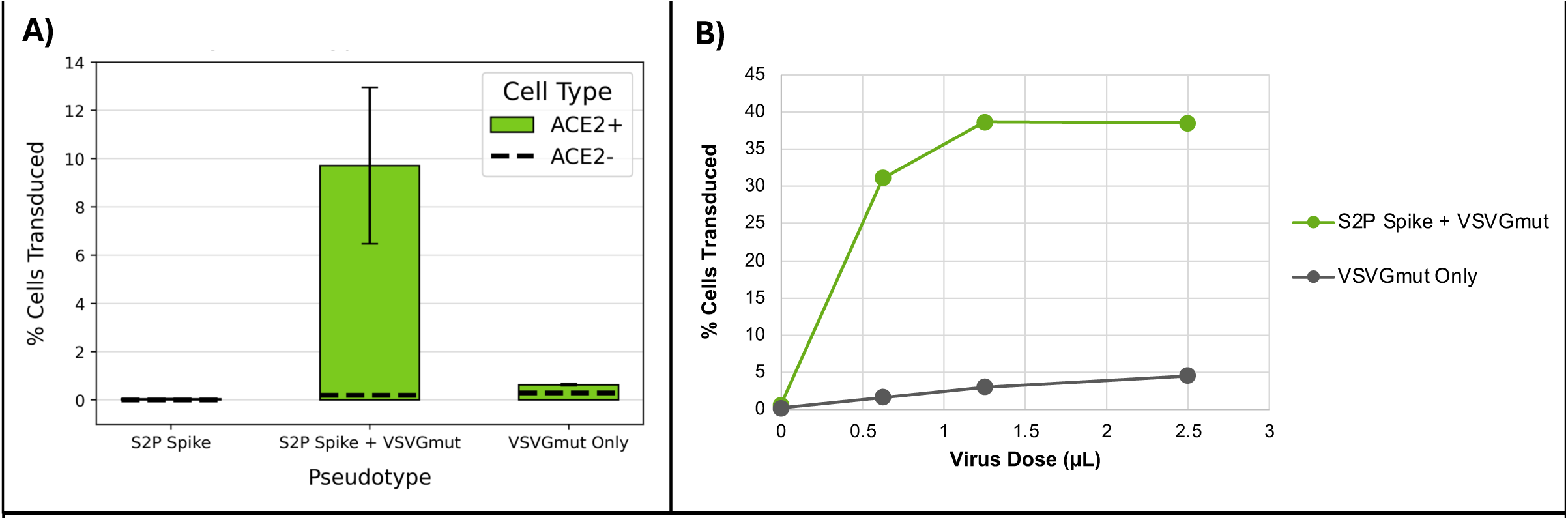
SARS-CoV-2 hybrid pseudotyped lentiviruses specifically infect ACE2-expressing cells. **A)** Transduction of ACE2+ and ACE2- cells by lentiviruses pseudotyped with either the Wuhan Strain SARS-CoV-2 prefusion-stabilized spike protein (S2P Spike), the fusion protein VSVGmut, or both. For each condition, 50 µL of unconcentrated lentivirus was added to fifty thousand cells. Transduction was measured via flow cytometry at 48 hours. Averages and standard deviations are plotted for the ACE2+ cell replicates (n = 5). Single replicates (n = 1) are plotted for the ACE2- cells. **B)** Transduction of ACE2+ cells at 48 hours by different doses of lentiviruses pseudotyped with both the Wuhan Strain SARS-CoV-2 S2P spike and VSVGmut (S2P Spike + VSVGmut) or VSVGmut alone (VSVGmut Only). Single replicates (n = 1) using concentrated virus and fifty thousand cells per well are plotted.

To convert the infection of ACE2-expressing cells into a neutralization assay, we mixed the viral protein-encoding lentiviral library with serum and measured the attenuation of infection in the presence vs absence of serum as a proxy for neutralization. We mixed CD and PPD sera with a VSVGmut-Wuhan Strain SARS-CoV-2 S2P spike lentivirus or the lentiviral library and incubated the mixtures with ACE2+ cells for 24 hours (**Figure 6A and 6B**). As with ARCADE, we observed that transduction rates in the neutralization assay were impacted by the serum dilutions used. CD sera reduced ACE2+ cell transduction by the lentiviruses in a dose-dependent manner, while highly diluted CD and PPD sera allowed transduction of ACE2+ cells at rates comparable to those allowed by the no serum controls. Given these data, dilutions from the more concentrated end of the range were used when performing neutralization assays.

**Figure 6.**
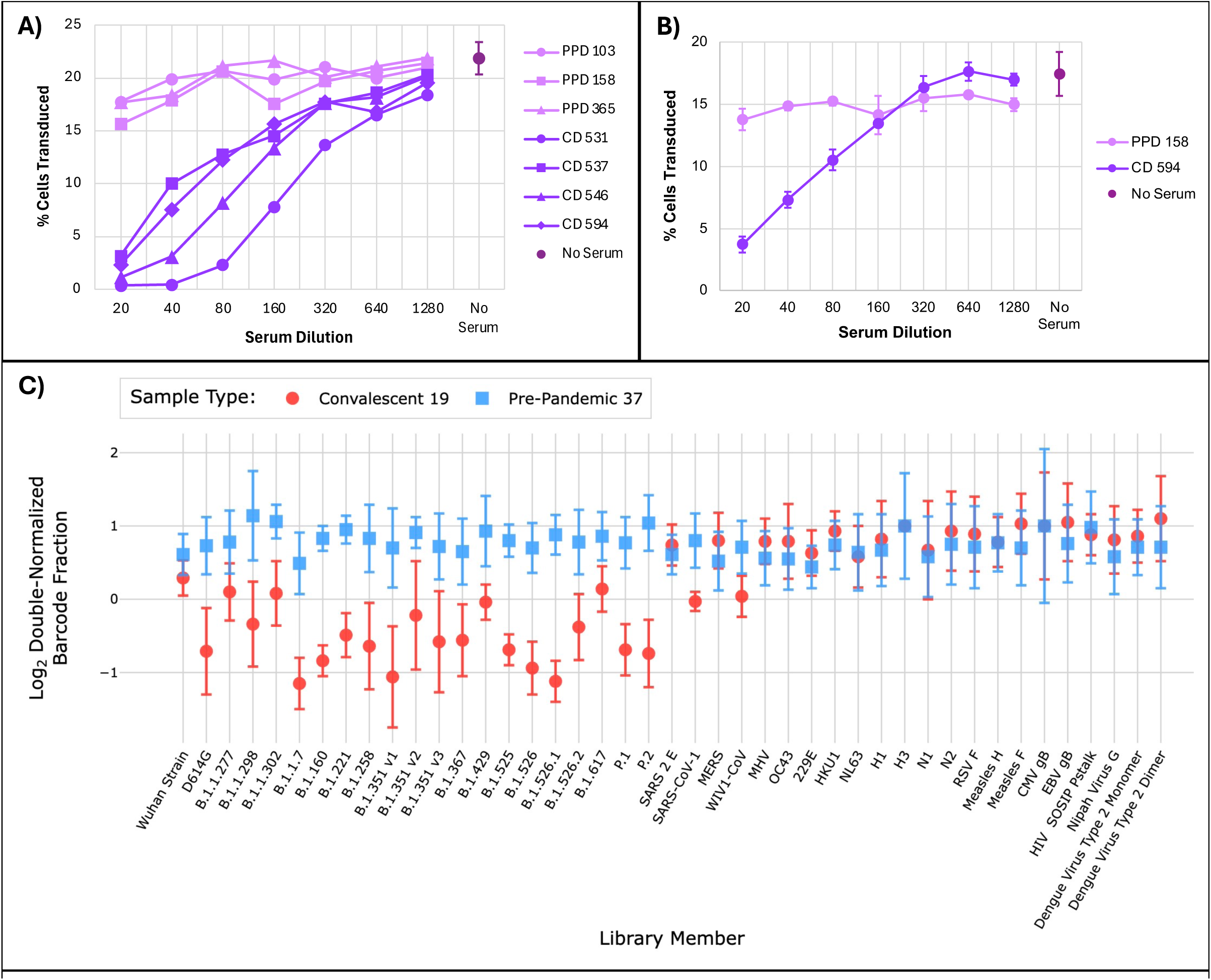
The neutralization assay differentiates between serum samples with and without SARS-CoV-2 neutralizing antibodies. **A)** Transduction of ACE2-expressing Jurkat cells by a lentivirus pseudotyped with the Wuhan Strain SARS-CoV-2 S2P spike and VSVGmut in the presence of serum at different dilutions. The lentivirus was mixed with convalescent donor (CD) or pre-pandemic donor (PPD) sera, and the mixtures were incubated with ACE2+ Jurkat cells. Cells were mixed with the lentivirus in the absence of serum as a control (No Serum). Flow cytometry was performed after 24 hours to measure transduction via cells’ expression of mCherry. Averages are plotted for CD and PPD samples (n = 2) and No Serum samples (n = 14). Error bars plotted for the No Serum samples correspond to the samples’ standard deviation. **B)** Transduction of ACE2-expressing Jurkat cells by the lentiviral library across serum dilutions. Library was mixed with CD or PPD sera, and the mixtures were incubated and analyzed as in **A**. Averages and standard deviations are plotted for CD and PPD serum samples (n = 3) and No Serum samples (n = 14). **C)** Double-normalized barcode fractions from a neutralization assay performed on CD and PPD sera. Sera were diluted 1:40, and the lentiviral library was diluted such that each sample received 0.5 µL of the library. The diluted sera and lentiviral library were mixed and incubated with cells for 48 hours. Transduction by each library member was measured via NGS of integrated barcodes. Averages and standard deviations of log_2_-transformed double-normalized barcode fractions are shown (n = 3). SARS 2 N CTD only and SARS 2 M had barcode fractions of zero for the No Serum samples, preventing double-normalization.

To apply our ARCADE pipeline to the neutralization assay, we first mixed 0.1, 0.5, or 1 µL of the lentiviral library with either media or serum diluted 1:40 or 1:80. Then, we incubated these mixtures with our ACE2+ cells for 48 hours and performed NGS. When analyzing double-normalized barcode fractions from the neutralization assay, we found that the barcode fractions for the SARS-CoV-2 variants, as well as for SARS-CoV-1 and WIV1-CoV, were generally smaller when the assay was performed on CD serum than when it was performed on PPD serum, showing the potential neutralization capacity of convalescent donors’ antibody repertoires (**Figure 6C**). While there was also detectable transduction of ACE2+ cells by control library members, including N1 and Measles H (**Supplementary Figure 11**), performing the assay with PPD and CD sera resulted in comparable barcode fractions, suggesting that this background signal was not neutralized by either serum type and was likely due to infection that occurred independently of ACE2, through other cell surface receptors or nonspecific endocytic mechanisms.^60–66^ These results suggest that our neutralization assay can be used to compare serum samples’ neutralization capacities across many viral antigens simultaneously.

## Discussion

Since emerging infectious diseases increasingly threaten human health,^3,27–30^ tools that can more efficiently and comprehensively characterize individuals’ and populations’ antibody-based immunity are required. ARCADE and multiplexed pseudovirus neutralization assays can survey the antigen recognition capabilities of serum antibodies in a high-throughput fashion to address this urgent need. By employing hybrid pseudotyped lentiviral libraries that link encoded barcodes to displayed antigens, our assays can characterize antibodies’ recognition of dozens of whole antigens in their native conformations simultaneously without the need for protein purification or immobilization. We have demonstrated that our assays identify the presence of antibodies against many SARS-CoV-2 variants concurrently and have validated that ARCADE’s sequencing readouts align with the antibody titers derived from established immunoassays.

We have also highlighted how ARCADE can facilitate better understanding of the protection conferred by vaccination and prior infections to different groups of individuals. We found that most individuals’ SARS-CoV-2 variant barcode fractions increased following vaccination but that the variants with the greatest increases tended to differ between the two COVID-19-vaccinated cohorts. Whereas most Moderna-vaccinated individuals saw the greatest increases in barcode fractions for variants from later in the pandemic (e.g., B.1.526.1), JCJ-vaccinated individuals saw the greatest increases for earlier variants like D614G and B.1.160, supporting studies showing that mRNA-based vaccines stimulate stronger and broader anti-SARS-CoV-2 antibody responses than adenovirus vector-based vaccines.^51–53^ Additionally, we found that both convalescent and vaccinated donors yielded low barcode fractions for the B.1.351 variants, reinforcing reports that these variants escaped existing humoral immunity.^45,51,54^ Given ARCADE’s ability to capture these key clinical findings, we anticipate that ARCADE could be readily applied to answer new questions about antibody responses in a range of infectious diseases.

While assays that employ lentiviruses and ACE2-expressing cell lines to characterize antibodies’ neutralization capabilities have been previously described,^17,18,67–70^ ARCADE enables large-scale measurements of antigen-specific antibodies even when an antigen-specific receptor cannot be expressed or remains unknown. This feature is particularly useful for measuring antibodies against emerging viruses, where knowledge of viruses’ antigenic proteins and the need for quantitative immunoprofiling assays may significantly outpace knowledge of viral entry receptors.

In this work, ARCADE employs a CD32-expressing cell line, which measures antibodies of the IgG isotype, with a preference for subclasses IgG1 and IgG3.^19,71–73^ Yet, other isotypes also play critical roles in a number of diseases.^74–77^ IgE, for example, serves as a crucial defense mechanism against parasitic infections; yet, it is also oftentimes responsible for harmful anaphylactic reactions.^78,79^ Exchanging CD32 in our cell line for other Fc receptors, such as FcεRI, could enable the study of a broader range of helpful and harmful antibody responses.

The highly modular nature of ARCADE enables it to also be applied outside the context of infectious disease. In principle, any protein that can be expressed on the surface of a lentiviral particle can be used for serum profiling. Thus, ARCADE could be adapted to study antibodies implicated in chronic diseases, such as cancer, autoimmunity and allergy.^78,80,81^ By pseudotyping lentiviruses with autoantigens, allergens, or other antigens, we could employ ARCADE to measure antibodies against these proteins, bolstering screening for a variety of immunological conditions and aiding the assessment of immunotherapies aimed at eliminating pathological antibody responses.

In conclusion, ARCADE can provide a wealth of data characterizing the antigens recognized by antibodies within human serum samples. By measuring antibodies’ relative reactivities to dozens of antigens simultaneously, ARCADE can enhance clinicians’ ability to screen patients for protective immunity and study the effects of diseases and clinical interventions on antibody responses.

## Limitations

While ARCADE seeks to improve the throughput and multiplexability of existing immunoassays, our current method of lentiviral library production limits the total number of antigens that can be surveyed at one time. In this work, we share results obtained with our forty-five-member lentiviral library as well as those from a proof-of-concept experiment suggesting that our assay could support multiplexed readouts obtained using lentiviral libraries with up to one hundred members (**Supplementary Figure 1A**). Performing deep mutational scans or assaying many variants of a wide range of viruses, however, would necessitate larger library sizes. Enhancing the antigen-specific transduction efficiency of our lentiviruses and reducing the complexity of library assembly, as recently described,^18^ could allow us to characterize antibody responses against the hundreds to thousands of antigens desired. Additionally, while lentiviral pseudotyping is a broadly generalizable technique, not all immunologically relevant proteins can be readily expressed on the surface of HEK producer cells, and therefore, cannot be properly displayed on the resulting lentiviral particles. The SARS-CoV-2 nucleocapsid protein may be one such protein, as our experiments did not count any barcodes for this library member, despite extensive evidence that both anti-nucleocapsid and anti-spike antibodies are generated during infection.^82–84^ We hypothesize that this lack of signal for the nucleocapsid protein is a consequence of expressing it outside of its native intraviral context.^84–87^ For proteins that do not typically localize to a cell’s plasma membrane, additional validation and optimization could be required for the proteins’ inclusion in our assay.

## Methods

### Ethics Statement

This work was reviewed and approved by the MIT Institutional Review Board (Protocol No. 2202000568).

### Media and Cells

HEK293T cells (ATCC CRL-11268) were cultured in DMEM (ATCC) supplemented with 10% fetal bovine serum (FBS; RCD Systems), penicillin-streptomycin (pen/strep; Gibco), and 25 mM HEPES (Rockland).

Jurkat cells (ATCC TIB-152) and K562 cells (ATCC CCL-243) were cultured in RPMI-1640 (ATCC) supplemented with 10% FBS and pen/strep.

### Plasmid Construction

The plasmids pMD2.G and psPAX2 were gifts from D. Trono (Addgene Plasmids #12259 and #12260, respectively). The pLentiCRISPR v2 plasmid was a gift from F. Zhang (Addgene Plasmid #52961). To generate the pLeAPS backbone, the CMV core promoter was cloned into the pLenti backbone between the polypurine tract and the 3′ LTR, analogous to previous work.^88^

For single lentivirus transductions, the Wuhan Strain SARS-CoV-2 prefusion-stabilized (S2P) spike was cloned into the pMD2 backbone. These transductions also utilized the VSVGmut-encoding plasmid available on Addgene (Plasmid #182229).

To assemble the forty-five-member lentiviral library, viral protein constructs described in **Supplementary Table 1** were cloned into the pLenti backbone and C-terminally fused to GFP via a P2A motif, and 8-nt barcodes were cloned into the pLeAPS backbone downstream of the Ef1α core promoter and mCherry. The pLeAPS backbone plasmid is available on Addgene (Plasmid #182230).

### Transfection for Lentiviral Production

Lentiviruses were produced by transient transfection of HEK293T cells with linear 25 kDa polyethylenimine (PEI; Santa Cruz Biotechnology) or *Trans*IT-Lenti (Mirus) at a 3:1 mass ratio of transfection reagent to DNA. Briefly, plasmids were diluted in Opti-MEM (Thermo Fisher) and mixed with transfection reagent to form complexes. Complex formation was allowed to proceed for ten to fifteen minutes at room temperature before dropwise addition to cells. If using PEI, the media was aspirated and replaced with fresh complete DMEM three to six hours post-transfection.

The plasmid mass ratios used for transfections were 5.6:3:3:1 for transfer plasmid to psPAX2 to envelope plasmid to fusogen plasmid. Not all components were used in each transfection. The transfer plasmids used were either pLenti (to produce viral antigen-displaying lentiviruses) or pLeAPS (to produce barcoded lentiviruses). Envelope plasmids were only used as an alternative to the pLenti plasmids, and each envelope plasmid consisted of a pMD2 backbone with a cassette that could express a viral antigen on the surface of the lentivirus. The fusogen plasmids used encoded either VSVG or VSVGmut.

### Lentiviral Harvest, Purification, and Concentration

Forty-eight hours post transfection, transfected HEK293T cell media containing unconcentrated lentivirus was collected and filtered through a 0.45-μm polyethersulfone filter (Millipore Sigma). If needed, the filtered virus was stored at 4°C for up to 2 weeks or at - 80°C indefinitely.

To concentrate the filtered lentiviruses by ∼200x, the viruses were subjected to ultracentrifugation at 100,000 x g for 90 minutes at 4°C. The supernatant was discarded, and lentivirus pellets were resuspended in Opti-MEM overnight at 4°C. Resuspended lentiviruses were aliquoted and stored at -80°C indefinitely.

### Generation of Lentiviral Libraries

To generate the forty-five-member lentiviral library, a packaging cell line was first produced. HEK293T cells were transfected as described above with psPAX2 and a plasmid encoding VSVG and either a pLeAPS mCherry-barcode-encoding plasmid or a pLenti GFP-viral antigen-encoding plasmid. After 48 hours, unconcentrated lentivirus was filtered and used to co-transduce HEK293T cells in duplicate in the presence of 8 µg/mL of either polybrene or diethylaminoethyl-dextran (Sigma-Aldrich). For these arrayed transductions, the cells were seeded at 20% confluency in 6-well plates (Thermo Fisher), and 500 µL of each of the corresponding barcode-encoding and viral antigen-encoding lentiviruses (**Supplementary Table 1**) were added per well. After 24 hours, the media was fully exchanged, and after 48 hours, cells were checked for transduction via flow cytometry. For each barcode-viral antigen pair, one duplicate was analyzed to determine the proportion of cells transduced with both antigen-encoding lentivirus (GFP+) and barcode-encoding lentivirus (mCherry+). Cells from the remaining duplicate were then pooled, using the proportion of double-positive cells to include an equal number of viral antigen-barcode-encoding cells for each member in the library. This pool of cells was then sorted based on GFP and mCherry expression to make the library packaging cell line.

To produce the forty-five-member lentiviral library from the packaging cell line, packaging cell line cells were transfected as described above using only psPAX2 and a VSVGmut-encoding plasmid. Both transfer and envelope plasmids were omitted from these transfections, as their components were accounted for by the barcode and viral antigen components encoded in the packaging cell line. After 48 hours, lentivirus was collected and concentrated as described above for use in our assays.

To generate ladder viruses, HEK293T cells were transfected with psPAX2, a VSVG-encoding plasmid, and a pLeAPS plasmid encoding a barcode not already present in the library. After 48 hours, lentiviruses were harvested and concentrated as described above.

Before combining the forty-five-member lentiviral library with the ladder viruses, titrations of each of the lentiviruses were performed on ACE2-expressing Jurkat cells. The total number of TUs in 1 µL of the library was used to calculate the numbers of TUs of ladder viruses to add. Ladder viruses with LeAPS barcodes 59, 67, and 83 were added at 10%, 1%, and 0.1% of the total TUs of the library, respectively. The complete library was aliquoted and stored at -80°C.

### Assay Cell Line Generation

The human ACE2 receptor and FcγRIIA (CD32) genes were each cloned into a pHIV backbone. Downstream of these genes, the backbone also contained an IRES followed by EGFR to provide a straightforward indicator of construct expression. These receptor-encoding plasmids, along with psPAX2 and a VSVG-encoding plasmid, were used to transfect HEK293T cells as described above. Lentivirus was harvested, filtered, concentrated, and used to transduce Jurkat cells. For the transductions, Jurkat cells were resuspended at a concentration of 1 million cells/mL and mixed with lentivirus at a 1:1 volume ratio in 6-well plates (1 mL of cells plus 1 mL of lentivirus). Polybrene was also added at a concentration of 8 µg/mL. Transduced Jurkat cells were sorted based on EGFR expression to establish the receptor-expressing cell lines.

### Serum Samples

Panels of serum samples were ordered from Access Biologicals. These included a panel of 15 donors vaccinated with the Moderna COVID-19 vaccine, a panel of 14 donors vaccinated with the JCJ COVID-19 vaccine, a panel of 87 donors that were negative for COVID-19, and a panel of 20 donors that were COVID-19 convalescent.

For the Moderna-vaccinated panel, samples had been collected between December 2020 and March 2021. Three samples were collected per donor. The first sample was collected 0– 7 days prior to the first vaccination (Pre-Vaccine), the second sample was collected 2–4 days prior to the second vaccination (Post Dose 1), and the third sample was collected 13–16 days after the second vaccination (Post Dose 2). Antibody titers were measured via the LIAISON SARS-CoV-2 S1/S2 IgG assay (Diasorin). The serum supplier stated that some subjects were known to have previous SARS-CoV-2 infections, but which donors had been infected was not specified. The donors had been tested and found negative for antibodies against HIV 1/2 and HCV and non-reactive for Hepatitis B surface antigen. Pooled samples had been found non-reactive for HIV-1 RNA, HBV DNA, and HCV RNA by a nucleic acid test.

For the JCJ-vaccinated panel, samples had been collected between March 2021 and September 2021. Three samples were collected per donor. The first sample was collected < 1 day prior to vaccination (Pre-Vaccine), the second sample was collected 13–14 days post vaccination (Two Weeks Post-Vaccine), and the third sample was collected 59–62 days post vaccination (Two Months Post-Vaccine). Antibody titers were measured via the LIAISON SARS-CoV-2 S1/S2 IgG assay (Diasorin). The serum supplier stated that some subjects were known to have previous SARS-CoV-2 infections, but which donors had been infected was not specified. The donors had been tested and found negative for antibodies against HIV 1/2 and HCV and non-reactive for Hepatitis B surface antigen. Pooled samples had been found non-reactive for HIV-1 RNA, HBV DNA, and HCV RNA by a nucleic acid test.

For the COVID-19 negative panel, samples had been collected between March 2017 and November 2018. These samples had been tested and found negative for antibodies against HIV 1/2 and HCV and non-reactive for Hepatitis B surface antigen. Pooled samples had been found non-reactive for HIV-1 RNA, HBV DNA, HCV RNA, WNV RNA, and Zika virus RNA by a nucleic acid test.

For the COVID-19 convalescent panel, samples had been collected in July 2020, and SARS-CoV-2 infection had been confirmed via a nasopharyngeal swab PCR test or another diagnostic method. Anti-SARS-CoV-2 antibody titers further confirming infection had also been obtained using the LIAISON SARS-CoV-2 S1/S2 IgG assay (Diasorin) and the ADVIA Centaur SARS-CoV-2 Total (IgG and IgM) assay (Siemens). The donors had been tested and found negative for antibodies against HIV 1/2 and HCV and non-reactive for Hepatitis B surface antigen. Pooled samples had been found non-reactive for HIV-1 RNA, HBV DNA, and HCV RNA by a nucleic acid test.

Additional pre-pandemic donor (103, 158, and 365) and convalescent donor (531, 537, 546, and 594) serum samples were obtained from Cellero (acquired by Charles River Laboratories).

Complete serum sample information for the vaccinated cohorts is available in **Supplementary Tables 2 and 3**.

### ARCADE

Serum samples were diluted in complete RPMI and mixed with concentrated lentivirus in 96-well plates. The lentivirus-serum mixtures were then incubated at 37°C for 30 minutes before 5 x 10^4^ CD32-expressing cells —K562 cells in the proof-of-concept experiments and CD32+ Jurkat cells in all other experiments — were added per well in a total assay volume of 100 µL. Following cell addition, plates were incubated at 37°C for 24 hours. Transduction rates were then quantified via flow cytometry or next-generation sequencing.

All noted serum dilutions correspond to the final dilution present in each assay well. For assays employing the lentiviral library, conditions were optimized to maintain representation across library members and, where indicated, ladder viruses were included to facilitate comparisons within and across samples.

### Pseudovirus Neutralization Assays

Serum samples were diluted in complete RPMI and mixed with lentivirus in 96-well plates. The lentivirus-serum mixtures were then incubated at 37°C for 30 minutes before 5 x 10^4^ ACE2-expressing Jurkat cells were added per well in a total assay volume of 100 µL. Following cell addition, plates were incubated at 37°C for 24–48 hours. Transduction rates were then quantified via flow cytometry or next-generation sequencing.

All noted serum dilutions correspond to the final dilution present in each assay well. For assays employing the lentiviral library, conditions were optimized to maintain representation across library members and, where indicated, ladder viruses were included to facilitate comparisons within and across samples.

### Flow Cytometry

Cells were resuspended in FACS buffer (PBS + 0.1% BSA and 1 mM EDTA) prior to performing flow cytometry on a Cytoflex S flow cytometer. Cell sorting was performed using the Sony MA900.

For each 96-well plate of ARCADE samples tested, three wells were devoted to flow cytometry controls: one Not Transduced (NT) well, which was provided only media in the place of lentivirus and serum, and two wells that were provided the lentiviral library and CD 594’s serum at a 1:160 dilution. Since the lentiviruses in our library integrate both mCherry and their library member barcode into transduced cells, we used flow cytometry to identify transduced assay cells by their expression of mCherry. If mCherry expression was observed in the CD serum wells and not the NT well, then we proceeded with gDNA extraction on all assay samples (see **Supplementary Figure 12** for our gating strategy and sample flow cytometry data).

### Genomic DNA (gDNA) Extraction and Next-Generation Sequencing (NGS)

Assay wells were mixed by pipetting, and their contents were transferred to V-bottom 96-well plates (Greiner). The plates were centrifuged at 300 x g for 5 minutes to pellet assay cells. The supernatant was decanted and 1x phosphate-buffered saline (PBS) was used to wash the cells. Centrifugation was repeated, and the supernatant was removed. If gDNA extraction postponement was necessary, cells were pelleted in a deep-well 96-well plate instead, and the plate was sealed and stored at -80°C until it was possible to proceed with gDNA extraction.

Pelleted cells were mixed with 6 µL of proteinase K (Revvity) and 200 µL lysis buffer (Revvity) and transferred to deep-well 96-well plates. The deep-well plates were sealed, vortexed for less than 5 seconds, and centrifuged at 300 x g for 1 minute at room temperature to bring all well contents to the bottom of each well. Plates were incubated in a 56°C-water bath for 10 minutes to promote protein digestion. The plate was again vortexed for less than 5 seconds and centrifuged at 300 x g for 1 minute at room temperature before being placed on ice.

The chemagic 360 instrument and chemagic DNA Tissue 10 Kit H96 (Revvity) were used to perform gDNA extraction. Integrated barcodes were then amplified via 30 cycles of PCR (see **Supplementary Table 4** for primer sequences). An additional 25 cycles of PCR were performed to attach index sequences and spacer nucleotides (**Supplementary Table 5**) as well as Singular’s adapter sequences to the ends of the PCR products. Following the second PCR, the samples were pooled by combining an equal volume of each sample. To purify the desired product, a double-sided size selection was performed on the pool using SPRIselect (Beckman Coulter) at a right ratio of 0.65x and a left ratio of 0.85x. When contaminating bands were still observed following the double-sided size selection, gel extraction and a left side size selection at a ratio of 2.5x were utilized to further purify the final product pool. The size-selected pool was sequenced on Singular’s G4 Sequencer using their 150 PE kit, and FASTQ files were prepared by MIT’s BioMicro Center.

### ELISAs

Anti-SARS-CoV-2 Antibody IgG Spike Trimer and Anti-SARS-CoV-2 (B.1.1.7) Antibody IgG Spike Trimer kits (ACROBiosystems) were used. ELISAs were performed according to kit instructions.

Serum antibodies are polyclonal in nature, meaning that they vary in their exact specificities, affinities, and abundances even if they recognize the same antigen. Due to these varying properties, ELISAs cannot provide exact titers for polyclonal antibody samples.^89^ Therefore, after performing these ELISAs, we reported the highest dilution at which the absorbance recorded was greater than 0.1. If no dilution resulted in an absorbance greater than 0.1, the titer was reported as zero or undetectable.

### Computational Analyses

All computational analyses were performed using Microsoft Excel or Jupyter Notebook 6.4.12 via Anaconda Navigator 2.3.2 with Python 3.9.13. Sample sizes were not predetermined using statistical methods.

The average Phred score was calculated for each read in each FASTQ file, and reads with scores less than 20 — less than 99% base call accuracy — were removed from the analysis pipeline. As barcodes within the library differed by a Hamming distance as little as two, high base call accuracy was essential for assigning reads to the correct library member (**Supplementary Figure 13**).

Barcodes were extracted from each read based on the barcode’s location in the read, and the extracted barcodes were compared to a dictionary of barcodes to update the barcode counts for each sample. Only exact barcode matches — with a Hamming distance of zero from a library member barcode — were counted. Double normalization was performed to enable barcode count comparisons within and across samples. First, a ladder virus with nonzero barcode counts for all samples was selected. This was the 1% ladder virus for ARCADE and the 10% ladder virus for the neutralization assay. For each sample, each library member’s barcode count was divided by the selected ladder virus’s barcode count for that sample. Then, to account for differences in baseline rates of transduction across the library, the ladder-normalized barcode fraction for each library member for each sample was divided by the average of the corresponding library member’s ladder-normalized barcode fractions for the no serum samples. Averages and standard deviations were then computed for each of these double-normalized barcode fractions. A pooled standard deviation was used when averaging the no serum samples, and error propagation was performed for all standard deviations to account for the divisions occurring in the normalization steps. Equations for these calculations are below, where X is any given member in the library and A is any given sample being assayed.

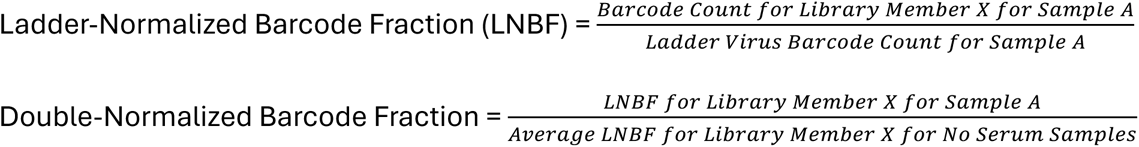

In some cases, normalizations, log_2_ transformations, or fold-change calculations were prevented due to certain library members having barcode fractions of zero. Library members for which this occurred were omitted from downstream analyses and visualizations. The average ladder-normalized barcode fraction for the library member SARS 2 N CTD only was zero for the no serum samples, preventing double normalization, and the library member SARS 2 M had double-normalized barcode fractions of zero for many samples, preventing other analyses.

Pearson correlations were performed using the linregress function from Python’s SciPy library, and principal component analysis (PCA) was performed using Python’s scikit-learn library. To perform PCA, features (double-normalized barcode fractions for each donor at all timepoints) were first transformed using StandardScaler. Then, PCA was performed using a range of numbers of components, and an elbow plot was assembled to assess the amount of variance explained by each component. K-means clustering was performed using the number of clusters determined by the Elbow Method.

Repeated-measures correlations were calculated using the rm_corr function from Python’s pingouin package. A mixed-effects linear model with a random intercept for each donor was fit using Python’s statsmodels library.

Figures were generated using Microsoft Excel or functions from Python’s Matplotlib, Seaborn, and Plotly libraries.

### Software

Plots were generated using Microsoft Excel and Jupyter Notebook. Flow cytometry data were analyzed by FlowJo (10.8.2).

## Acknowledgements

We thank the Koch Institute’s Flow Cytometry Core and the MIT BioMicro Center for their technical support. We thank S. Levine and C. Hallee for many helpful discussions and suggestions regarding next-generation sequencing. We thank A. Toniappa for providing training in using the chemagic 360 instrument and significant assistance in implementing our gDNA extraction protocol on the instrument.

This work was supported by Department of Defense Award W81XWH-22-1-0300 to M.E.B., a National Science Foundation Graduate Research Fellowship to A.C.H., a National Science Foundation Graduate Research Fellowship and a Siebel Scholarship to C.S.D., and a Canadian Institutes for Health Research Doctoral Foreign Study award to S.A.G. Core facilities in the Koch Institute are partially supported by Cancer Center Support (core) grant P30-CA014051 from the National Cancer Institute. This research is additionally supported by the National Research Foundation, Prime Minister′s Office, Singapore under its Campus for Research Excellence and Technological Enterprise program, through Singapore MIT Alliance for Research and Technology: Critical Analytics for Manufacturing Personalized-Medicine Inter-Disciplinary Research Group.

This material is based upon work supported by the National Science Foundation Graduate Research Fellowship Program under Grant No(s) 1122374, 1745302, and 2141064. Any opinions, findings, and conclusions or recommendations expressed in this material are those of the author(s) and do not necessarily reflect the views of the National Science Foundation.

## Author Contributions

C.S.D. and M.E.B. devised ARCADE. C.S.D. constructed the forty-five-member lentiviral library packaging cell line. C.S.D. and L.C.W. conducted and analyzed data for proof-of-concept experiments and produced the assay cell lines. M.E.B., C.S.D., and L.C.W. devised the data normalization strategy. L.C.W. obtained the panels of serum samples. S.A.G. contributed to early library troubleshooting. A.C.H. and L.C.W. generated the complete library with ladder viruses. A.C.H. conducted all assays and analyses pertaining to the vaccinated cohort samples, performed and analyzed data for the ELISAs, and performed and analyzed data for all ARCADE and neutralization assays leading to NGS readouts. M.E.B. supervised the research. A.C.H. wrote the manuscript. All authors contributed to the review and editing of the manuscript.

## Data Availability

The next-generation sequencing datasets generated and analyzed for this work are available in the National Center for Biotechnology Information Sequence Read Archive (BioProject Accession: PRJNA1364758).

## Code Availability

A sample Python Jupyter notebook for extracting and normalizing library member barcode counts is available in the Birnbaum Lab’s code repository: https://github.com/birnbaumlab/ARCADE

