## Supplemental information for "Interrogating antiviral antibody responses with multiplexed, high-throughput serum assays"

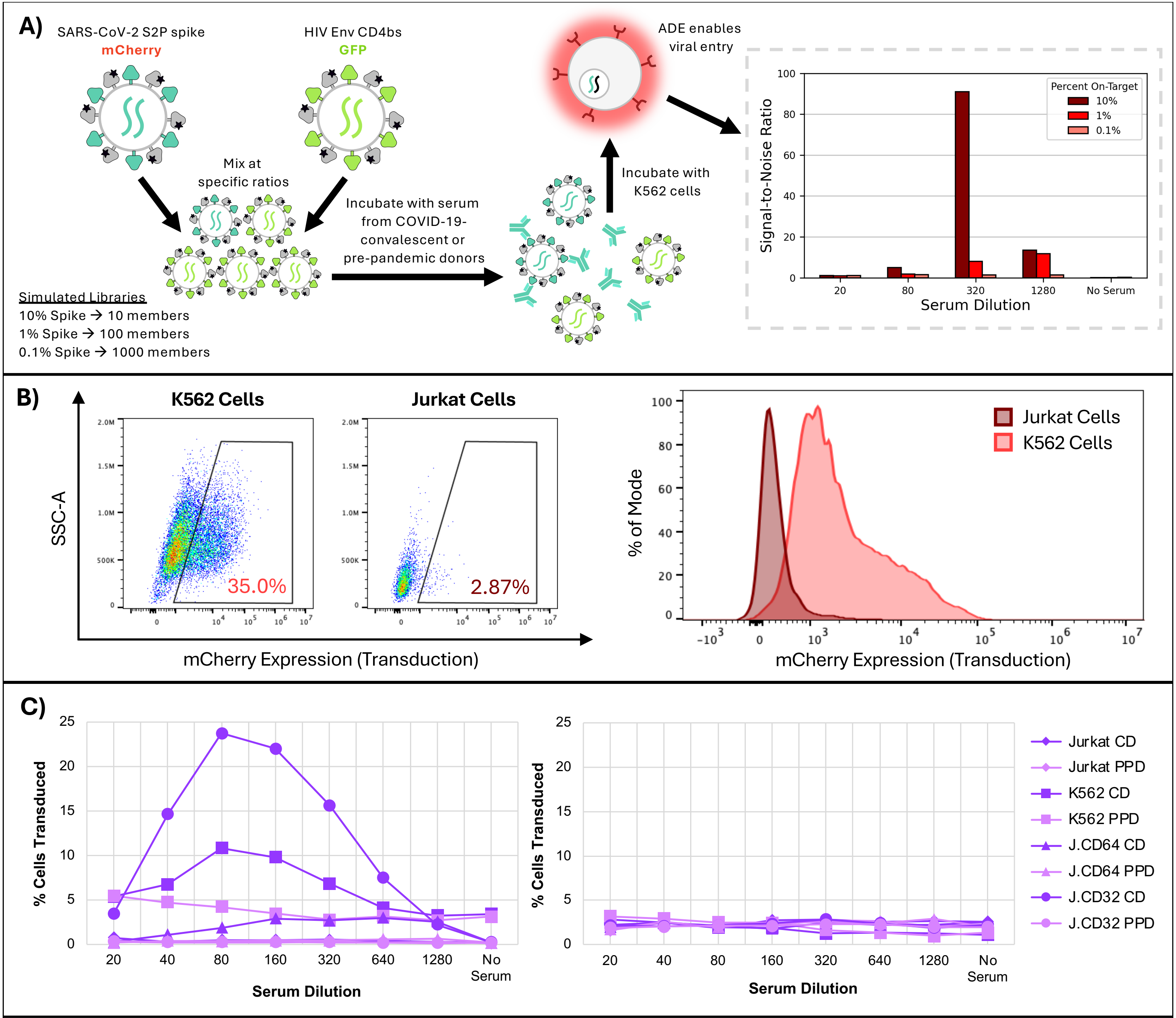


**Supplementary Figure 1.** ARCADE optimizations. **A)** Signal-to-noise ratios from simulating lentiviral libraries of different sizes. Lentiviruses were hybrid pseudotyped with VSVGmut and either on-target Wuhan Strain SARS-CoV-2 S2P spike or off-target HIV envelope glycoprotein, specifically the CD4 binding site from gp120/gp41. On-target lentiviruses, encoding mCherry, were combined with off-target lentiviruses, encoding GFP, at ratios simulating lentiviral libraries of 10, 100, or 1000 members, and the mixtures were incubated with pre-pandemic donor (PPD) 158 or COVID-19-convalescent donor (CD) 594 serum and K562 cells, which express the IgG-binding Fc receptor CD32. Flow cytometry determined the percentage of cells transduced by each lentivirus. Signal-to-noise ratios (S:N) for each simulated library were calculated using the following equations: Signal-to-Noise Ratio = (CD serum enrichment) / (PPD serum enrichment), where enrichment = (% mCherry+ / % GFP+) / (starting frequency of on-target virus). Single replicates (n = 1) are plotted. **B)** Background transduction of K562 cells and Jurkat cells by the lentiviral library. Fifty thousand K562 or Jurkat cells were mixed with the lentiviral library in the absence of serum. Transduction (mCherry expression) was measured at 24 hours via flow cytometry. Single representative replicates (n = 1) are shown. **C)** On-target (SARS-CoV-2 S2P spike, left) and off-target (HIV envelope glycoprotein, right) transduction of different cell lines in the presence of PPD 158 or CD 594 serum. The on-target and off-target lentiviruses were mixed 1:1 by volume, and the mixture was added to each assay well. Single assay replicates (n = 1) are shown. Jurkat cells express no Fc receptors, while K562 cells endogenously express FcγRII (CD32).[33] Jurkat cells were also engineered to express FcγRI (J.CD64) or FcγRIIA (J.CD32).

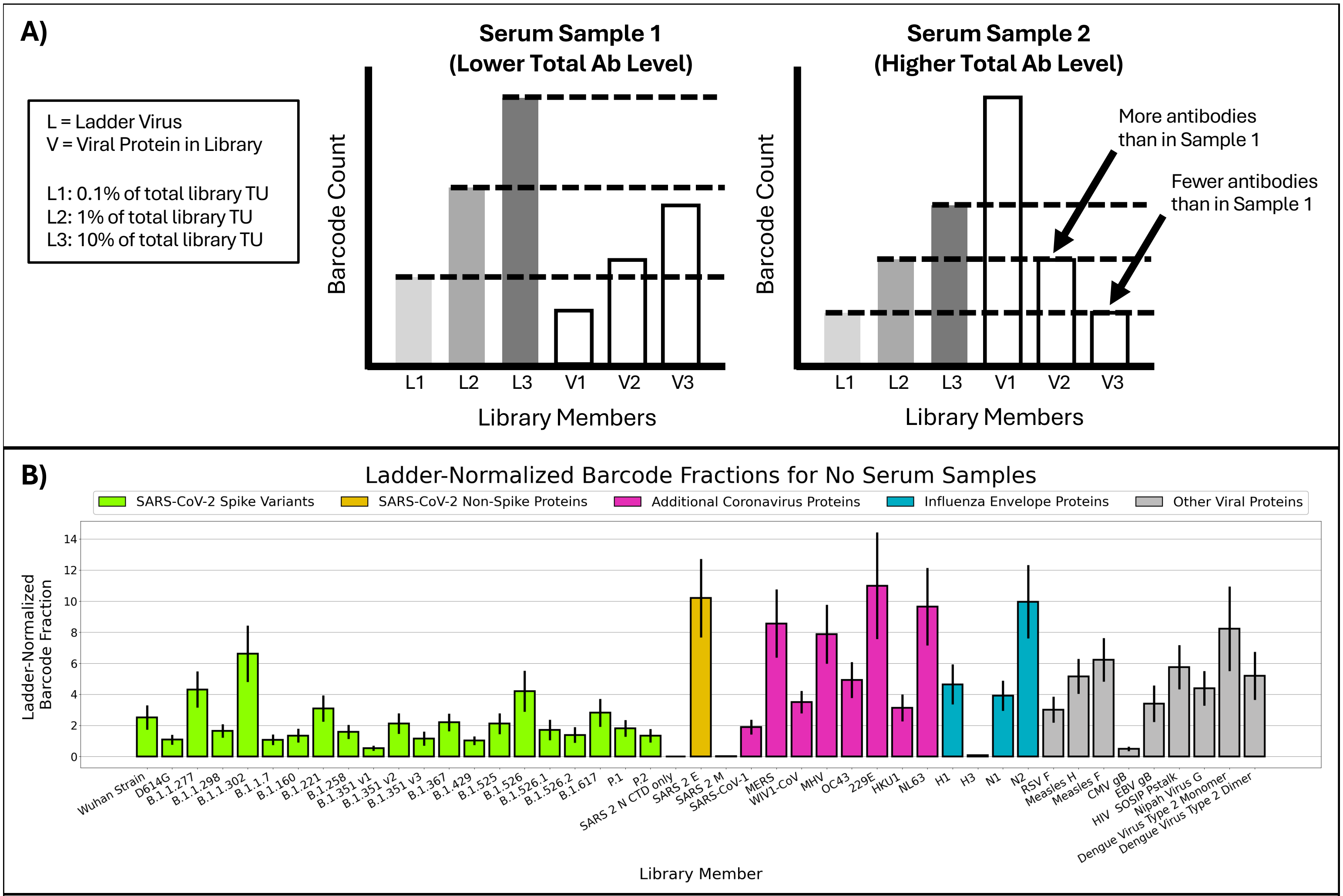


**Supplementary Figure 2.** Strategy for double-normalizing assay barcode fractions. **A)** Mock data demonstrating the utility of ladder viruses when comparing ARCADE barcode counts for certain library members across samples. The ladder viruses (L1, L2, and L3) transduce all cells at equal rates via the LDL receptor. These transduction rates are ideally unaffected by the presence, absence, or composition of the serum used in the assay. Individual ladder viruses encoding different barcodes are added to the library in amounts equivalent to 0.1%, 1%, and 10% of the library's total transducing units (TU). Adding these ladder viruses to the library allows the relative amounts of library member recognizing-antibodies to be compared across samples. **B)** Average ARCADE 1% ladder virus-normalized barcode fractions for No Serum samples across the lentiviral library. Each bar represents twenty-one technical replicates (n = 21), three replicates for each of seven assay plates. Error bars represent standard deviations. The average ladder virus-normalized barcode fraction for library member "SARS 2 N CTD only" is zero.

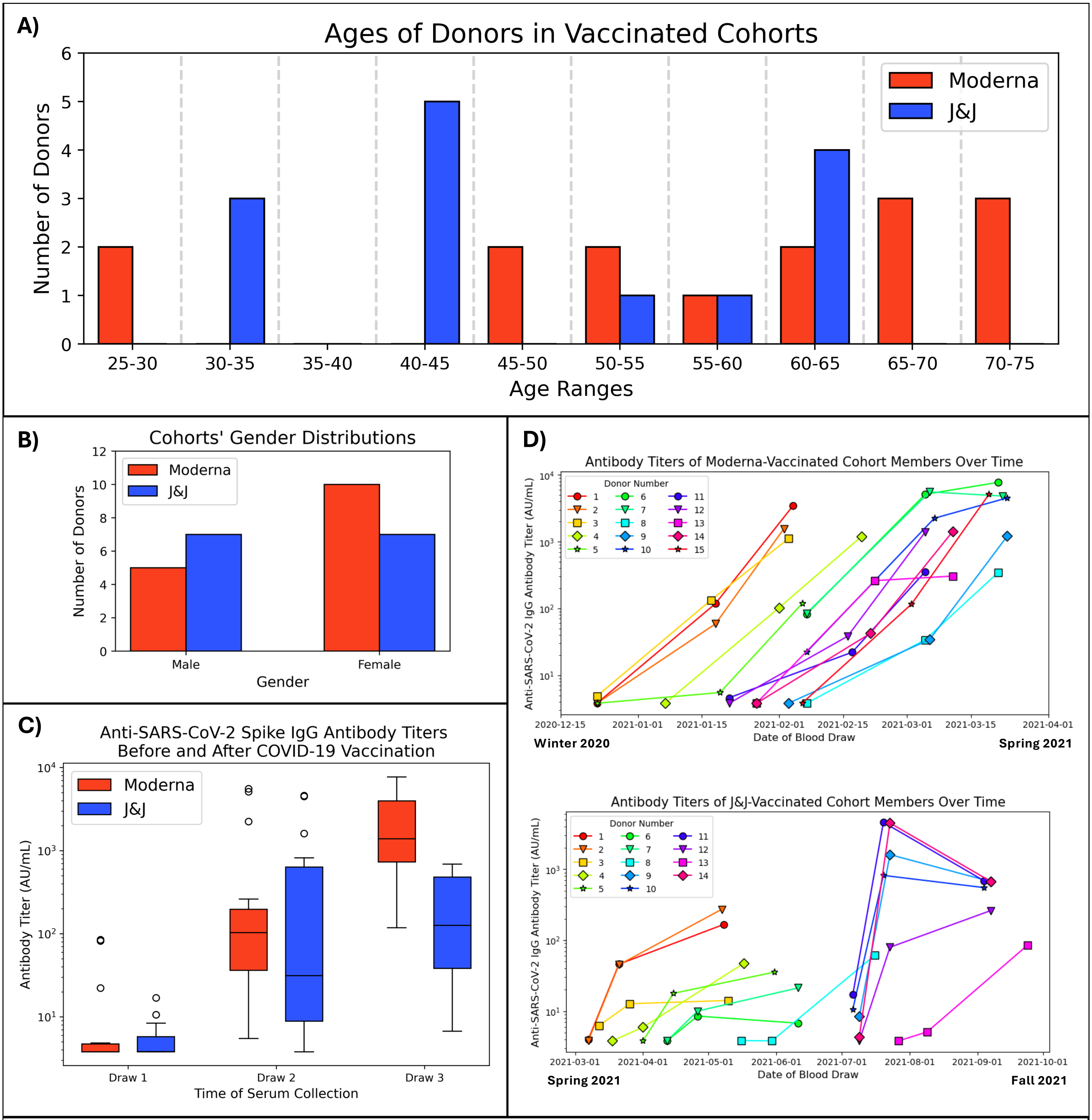


**Supplementary Figure 3.** Demographic information for donors from the vaccinated cohorts. **A)** Ages of donors by cohort. **B)** Genders of donors by cohort. **C)** Distributions of anti-SARS-CoV-2 S1/S2 IgG antibody titers for each timepoint by cohort. Titers were provided by the serum supplier and were measured via a LIAISON assay. Draw 1 = Pre-Vaccine, Draw 2 = Post Dose 1 (Moderna) or Two Weeks Post-Vaccine (J&J), Draw 3 = Post Dose 2 (Moderna) or Two Months Post-Vaccine (J&J). **D)** Antibody titers of donors over time.

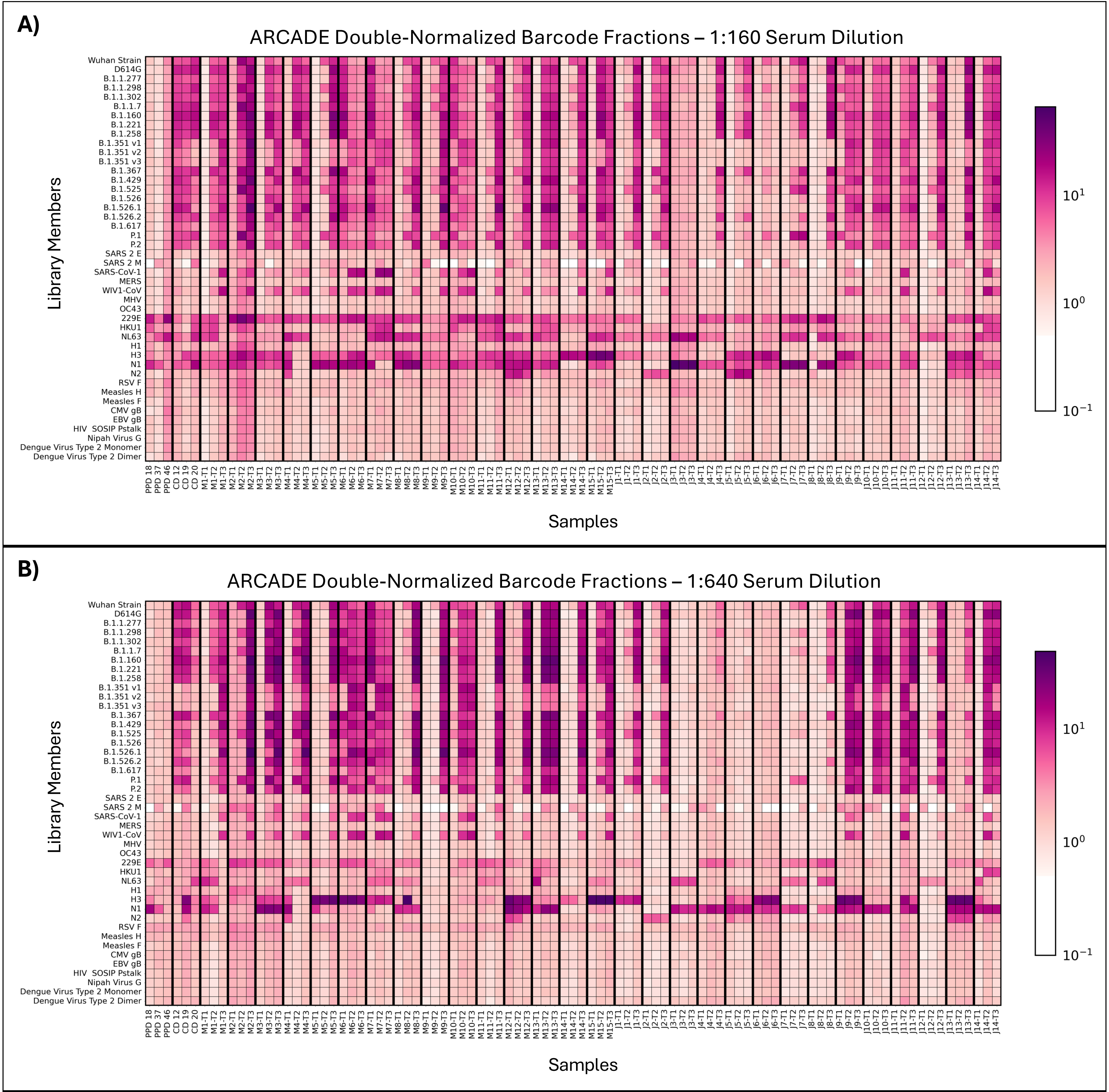


**Supplementary Figure 4.** Double-normalized barcode fractions from ARCADE performed with the lentiviral library on serum from pre-pandemic donors (PPD), convalescent donors (CD), and vaccinated donors. Double-normalized barcode fractions for each serum sample were averaged (n = 3). Sera were diluted 1:160 (**A**) or 1:640 (**B**). M = Moderna, J = J&J; T1 = Pre-Vaccine, T2 = Post Dose 1 (Moderna) or Two Weeks Post-Vaccine (J&J), T3 = Post Dose 2 (Moderna) or Two Months Post-Vaccine (J&J).

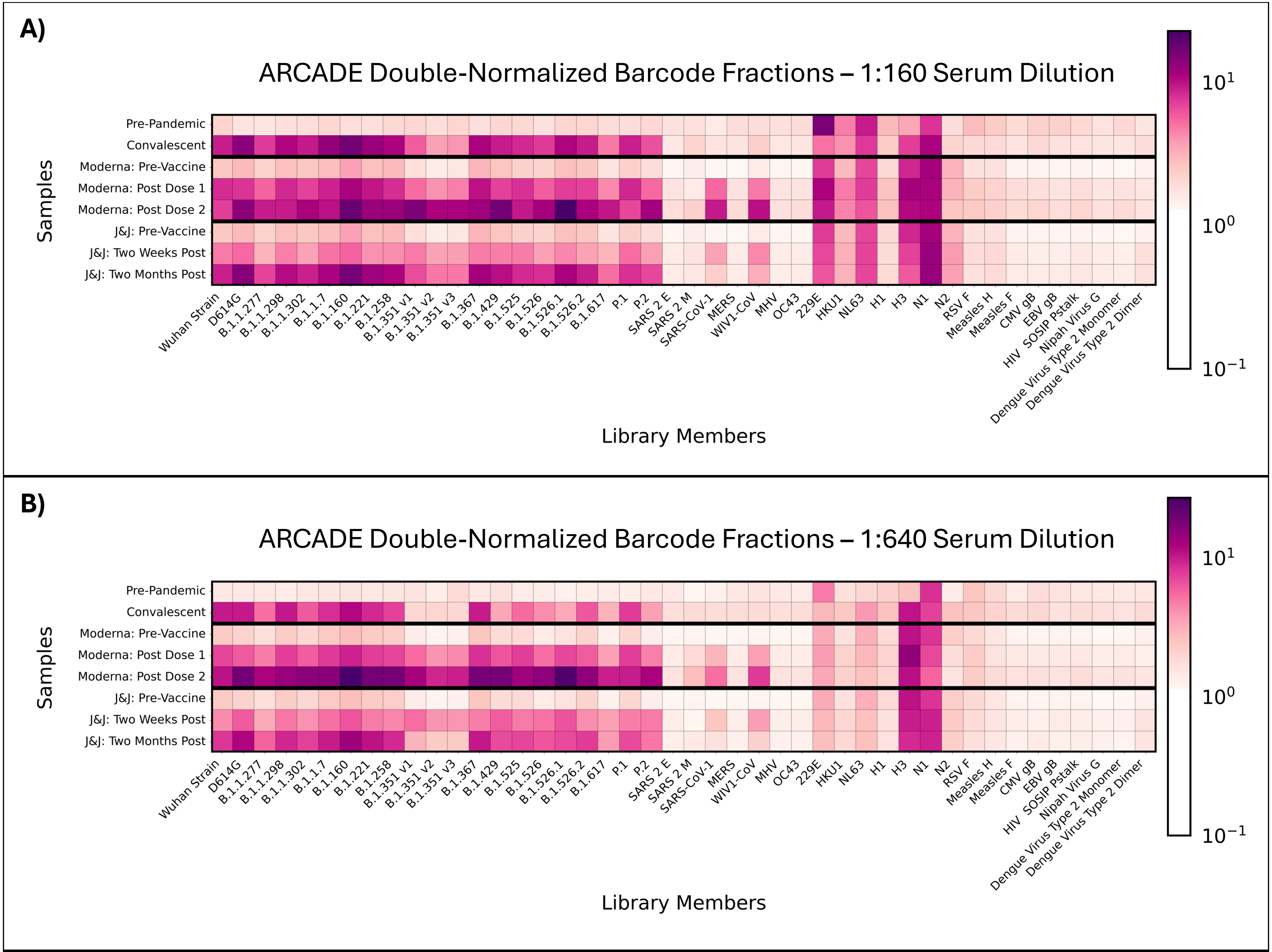


**Supplementary Figure 5.** Double-normalized barcode fractions for serum samples grouped by condition. ARCADE was performed using the lentiviral library and sera from individuals vaccinated with either the Moderna or J&J COVID-19 vaccine as well as sera from COVID-19-convalescent individuals and sera that were collected from healthy individuals pre-pandemic. Serum dilutions of 1:160 and 1:640 were tested. Barcode fractions were normalized using the 1% ladder virus, averaged across replicates (n = 3) and donors (Moderna: n = 15, J&J: n = 14, Pre-Pandemic: n = 4, Convalescent: n = 4), and further normalized using No Serum sample barcode fractions. The pre-pandemic and convalescent donors are used as negative and positive controls, respectively. **A**) Barcode fractions at the 1:160 dilution. **B)** Barcode fractions at the 1:640 dilution.

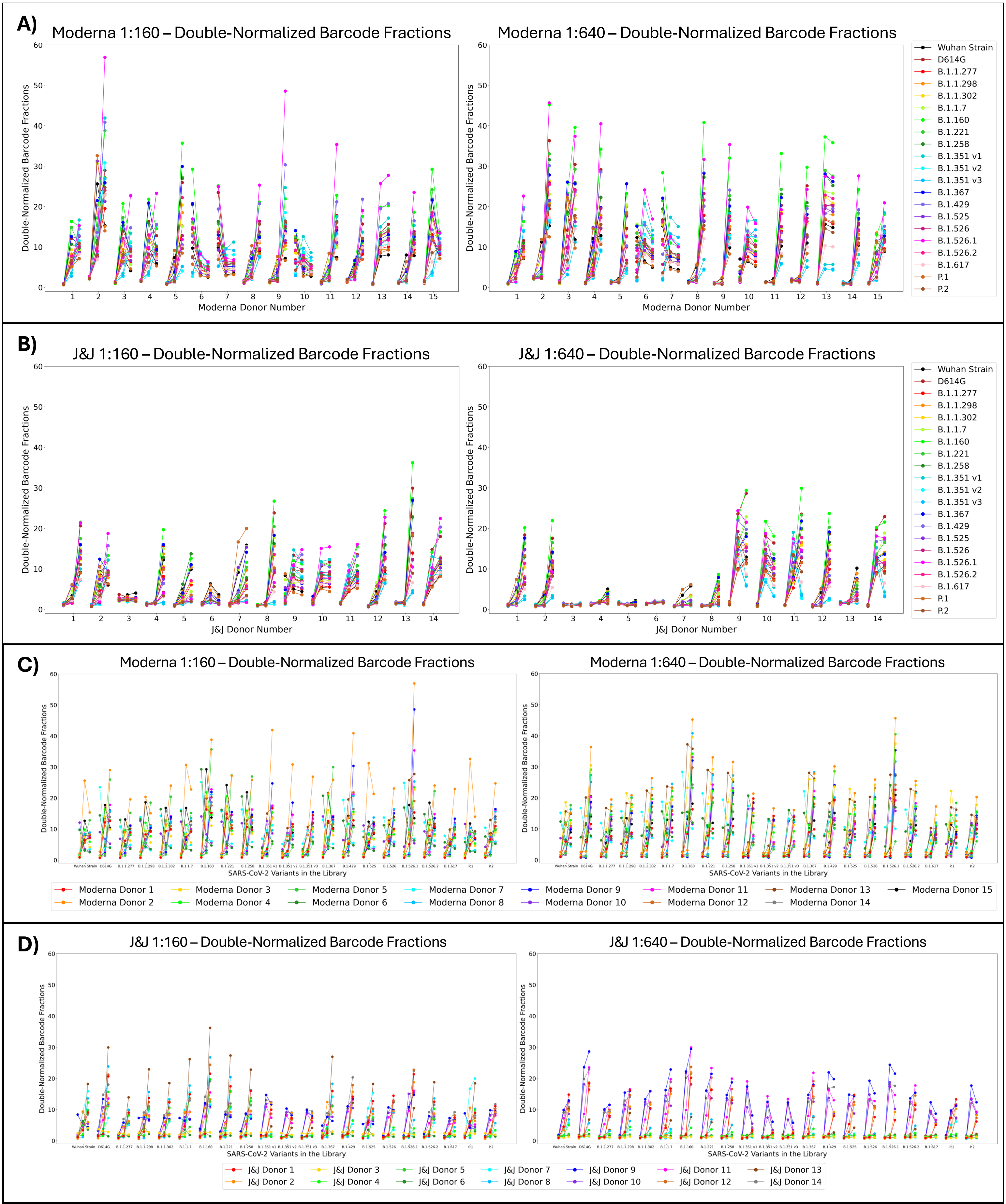


**Supplementary Figure 6.** Individual COVID-19 vaccine-induced anti-SARS-CoV-2 antibody responses characterized by ARCADE using the lentiviral library. ARCADE was performed on sera from individuals vaccinated with either the Moderna or J&J COVID-19 vaccine. Barcode fractions were normalized using the 1% ladder virus, averaged across replicates (n = 3), and further normalized using No Serum samples' barcode fractions. **A)** and **C)** highlight the Moderna-vaccinated cohort samples at dilutions 1:160 and 1:640, respectively, while **B)** and **D**) highlight the J&J-vaccinated cohort samples at dilutions 1:160 and 1:640, respectively. **A)** and **B)** highlight differences between donors, whereas **C)** and **D)** highlight differences between the SARS-CoV-2 variants in the library.

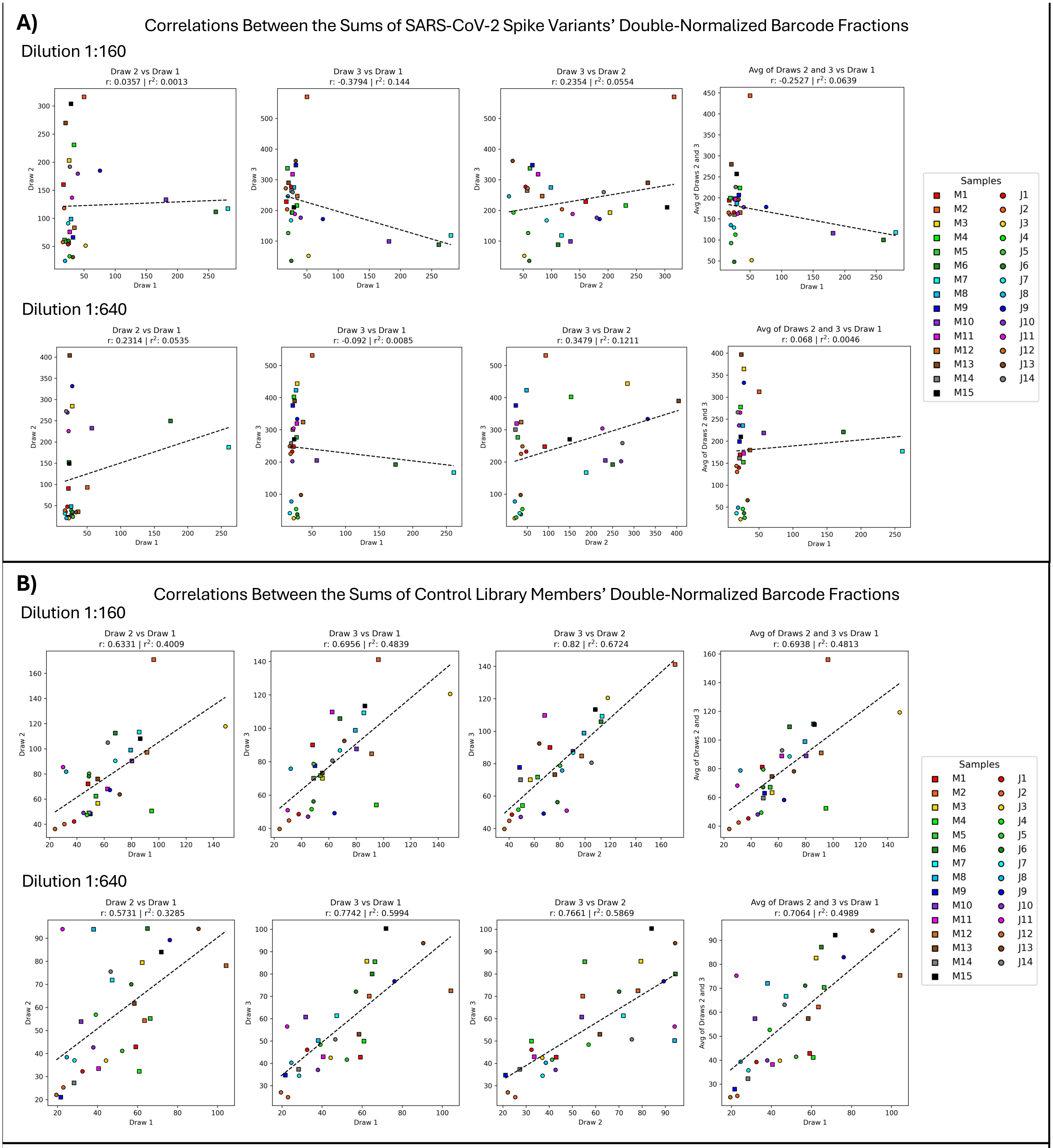


**Supplementary Figure 7.** Correlations between vaccinated donors' double-normalized barcode fractions across blood draws. **A)** Correlations between barcode fractions for SARS-CoV-2 spike variants pre- and post-vaccination. For each timepoint for each donor, double-normalized barcode fractions for all SARS-CoV-2 spike variants in the library were summed. Pearson correlation coefficients were calculated to describe the relationships between barcode fractions from different blood draws. **B)** Correlations between barcode fractions for non-SARS-CoV-2 library members pre- and post-vaccination. For each timepoint for each donor, double-normalized barcode fractions for all non-SARS-CoV-2 library members were summed. Pearson correlation coefficients were calculated to describe the relationships between barcode fractions from different blood draws. M = Moderna, J = J&J; Draw 1 = Pre-Vaccine, Draw 2 = Post Dose 1 (Moderna) or Two Weeks Post (J&J), Draw 3 = Post Dose 2 (Moderna) or Two Months Post (J&J), Draw 2/3 = Average of Draw 2 and Draw 3.

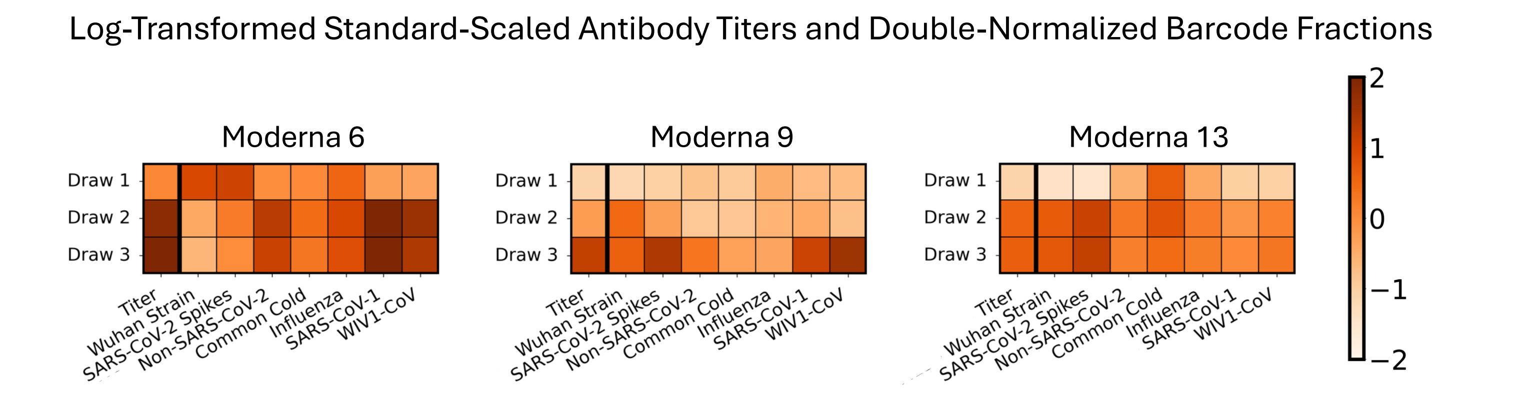


**Supplementary Figure 8.** Examples of alignment between anti-SARS-CoV-2 antibody titers and SARS-CoV-1's and WIV1-CoV's barcode fractions as compared to other library members' barcode fractions. Antibody titers were provided by the serum supplier and had been measured with a LIAISON SARS-CoV-2 S1/S2 IgG assay. Titers and average (n = 3) double-normalized barcode fractions — from ARCADE performed with sera diluted 1:640 — underwent a $\log_{10}(x + 1)$ transformation followed by standard scaling. Titers were scaled across all donors and timepoints, while double-normalized barcode fractions were scaled within their barcode fraction type across all donors and timepoints. The labels SARS-CoV-2 Spikes, Non-SARS-CoV-2, Common Cold, and Influenza refer to sums of barcode fractions for library members with barcode numbers 1-10 and 18-28; 11-17, 29, and 33-45; 13-16; and 33-36, respectively. A sampling of donors who were vaccinated with the Moderna COVID-19 vaccine is shown. Draw 1 = Pre-Vaccine, Draw 2 = Post Dose 1, Draw 3 = Post Dose 2.

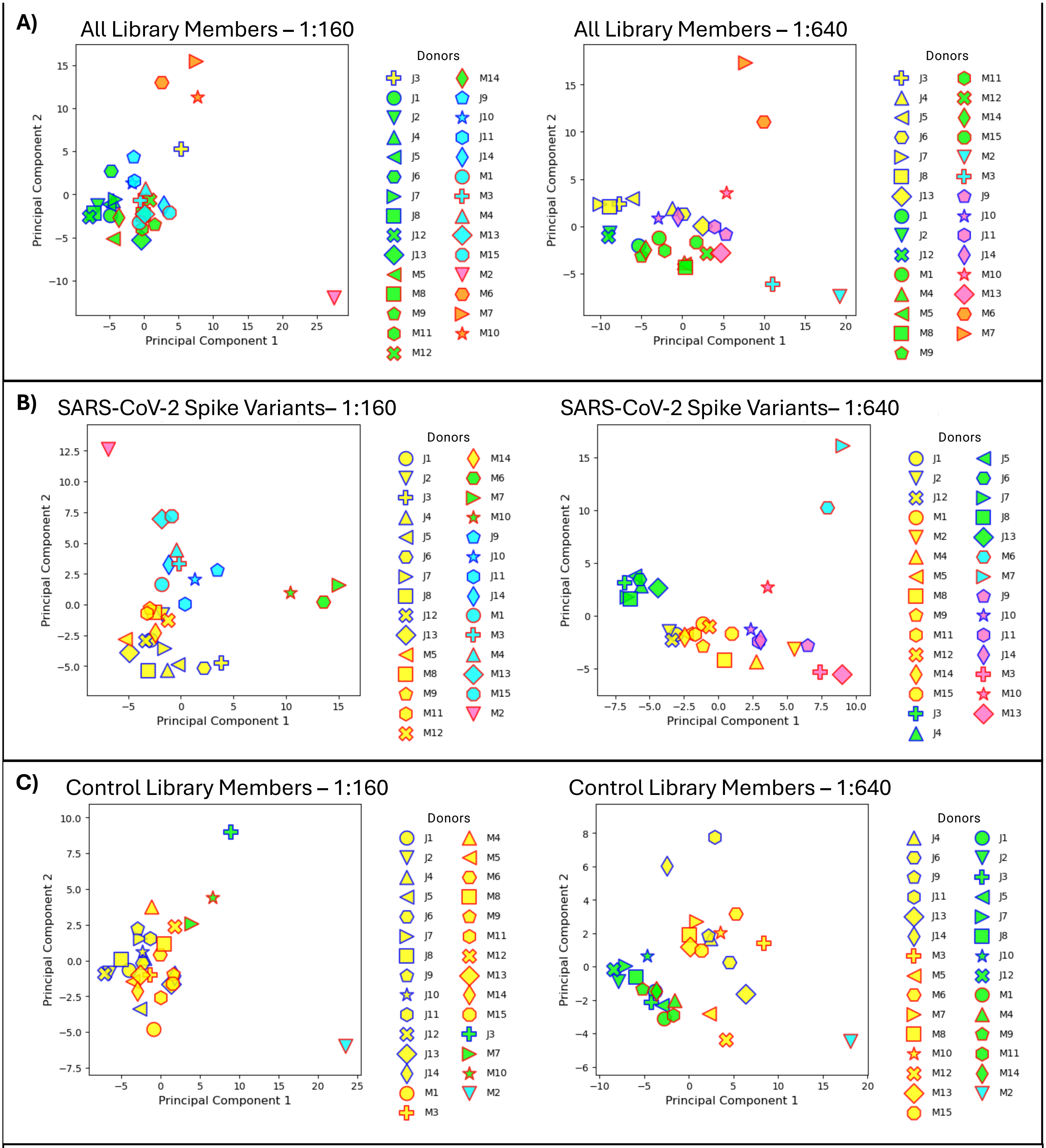


**Supplementary Figure 9.** Clustering of vaccinated donors based on their double-normalized barcode fractions from ARCADE. Principal component analyses (PCA) were performed with each feature consisting of a donor's set of double-normalized barcode fractions for each library member at each timepoint. Each point in principal component space corresponds to a donor vaccinated with the Moderna (M) or J&J (J) COVID-19 vaccine. Results from two serum dilutions (1:160 (left) and 1:640 (right)) used in ARCADE are shown. PCA were repeated with barcode fractions from different subsets of the library: all library members (**A**), SARS-CoV-2 spike variants only (**B**), and control library members only (**C**). K-means clustering was performed with 5 (**A**), 4 (**B**), or 3 (**C**) clusters based on the Elbow method. The interior color of each point indicates its cluster identity, and the outline color of each point indicates whether the corresponding donor received the Moderna (red) or J&J (blue) vaccine.

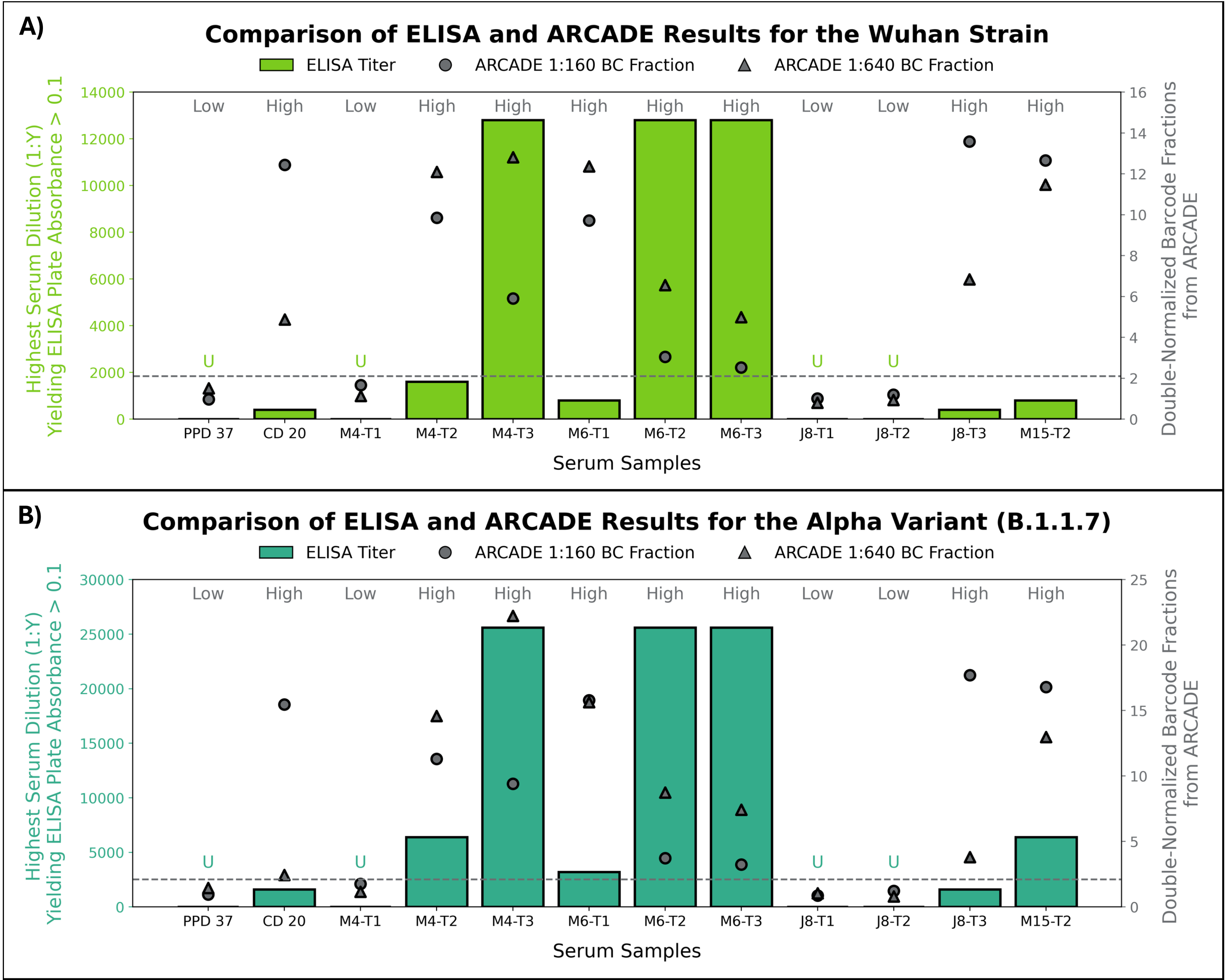


**Supplementary Figure 10.** Alignment between double-normalized barcode fractions from ARCADE using the lentiviral library and antibody titers from ELISAs. Anti-SARS-CoV-2 spike trimer IgG ELISAs were performed to measure antibodies against two different SARS-CoV-2 library members, the wildtype Wuhan Strain (**A**) and the Alpha variant (B.1.1.7) (**B**). Each ELISA was performed on samples that varied in their SARS-CoV-2 barcode fractions. Samples with undetectable antibody levels via ELISA — those with absorbances less than 0.1 for all tested dilutions — are labeled "U". Dotted lines demarcate barcode fraction thresholds (**A**: 2.10; **B**: 2.11), above which samples tend to have detectable antibody titers via ELISAs. Samples are classified as containing "Low" or "High" levels of anti-SARS-CoV-2 antibodies based on the detectability of those antibodies via ELISA and ARCADE.

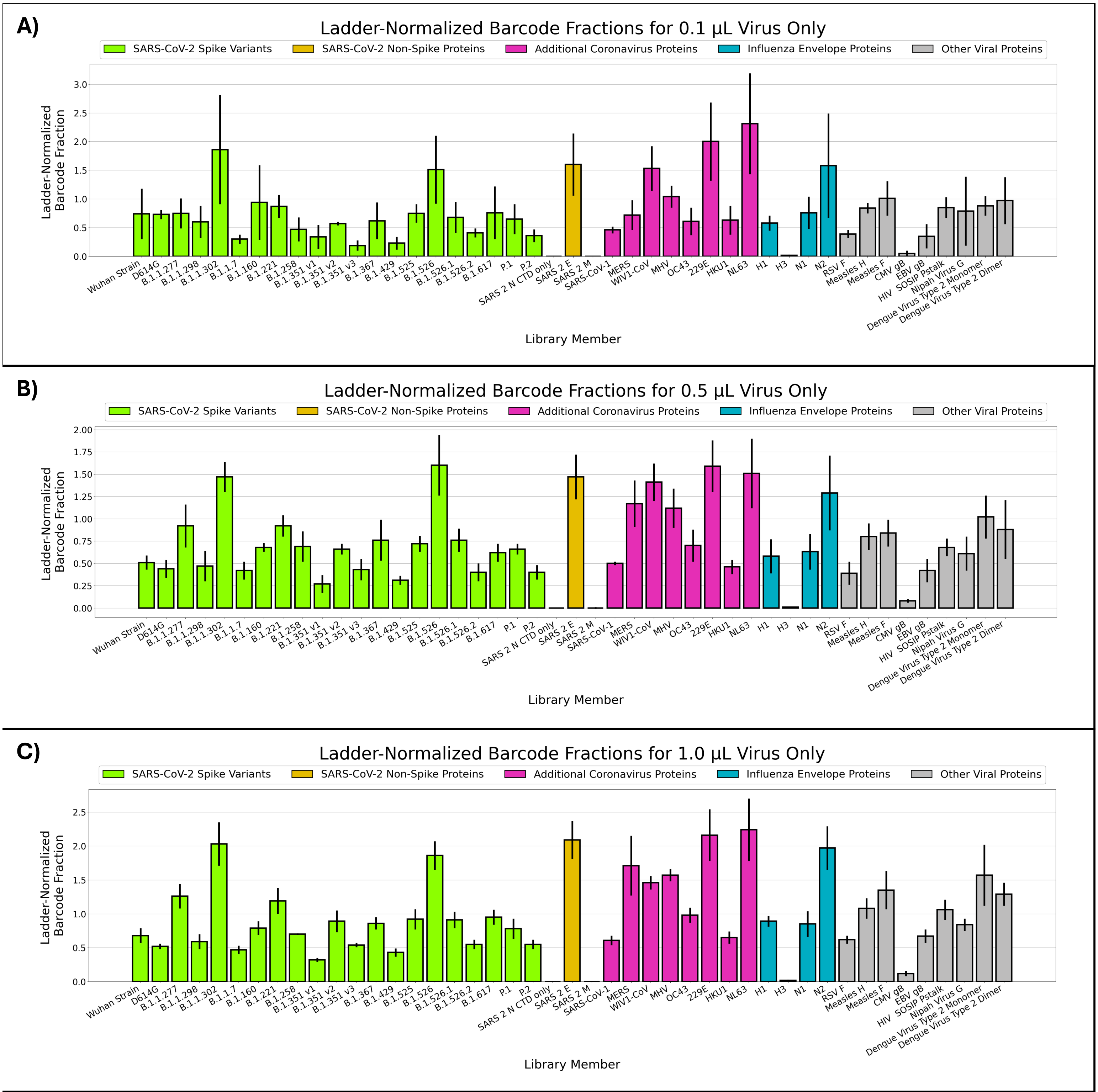


**Supplementary Figure 11.** Average 10% ladder virus-normalized barcode fractions for No Serum samples in the neutralization assay. The lentiviral library was added to ACE2-expressing cells in volumes equivalent to 0.1 µL (**A**), 0.5 µL (**B**), or 1.0 µL (**C**). After a 48-hour incubation, gDNA was extracted from assay cells, and integrated library member barcodes were amplified in preparation for next-generation sequencing. Each sample's library member barcode fractions were divided by the sample's 10% ladder virus barcode fraction. Averages with standard deviations (n = 3) are plotted. The average ladder virus-normalized barcode fractions for library members "SARS 2 N CTD only" and "SARS 2 M" are zero.

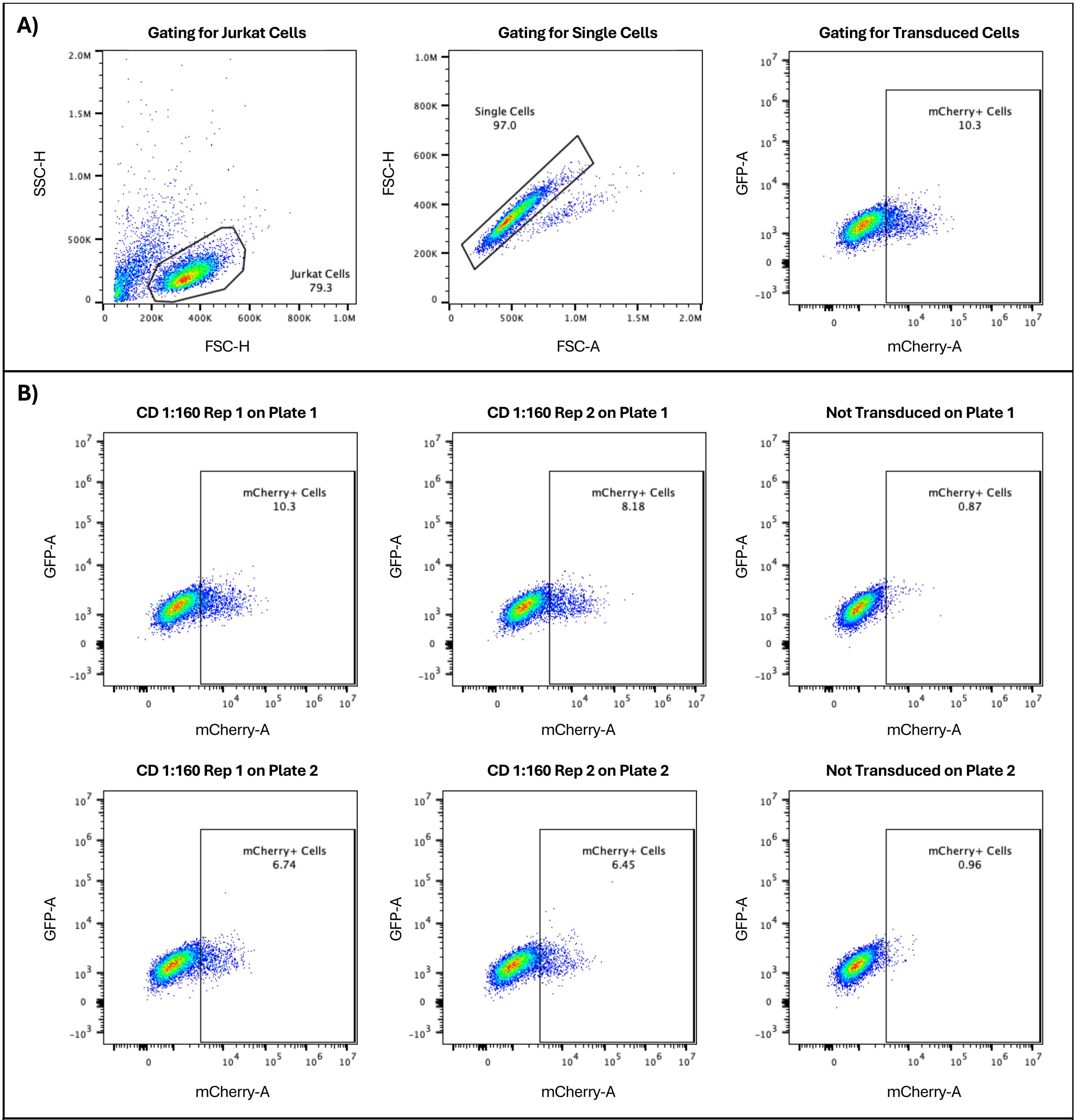


**Supplementary Figure 12.** Representative flow cytometry data from ARCADE using the lentiviral library. **A)** Gating strategy used to measure assay cell line transduction rates. **B)** Flow cytometry data for ARCADE control samples. Convalescent donor (CD) serum samples diluted 1:160 resulted in transduction rates of 5-10%, as determined by mCherry signal. For assay wells that received cells but not lentivirus (Not Transduced), less than 1% of cells were mCherry+. Control samples from two different assay plates are shown.

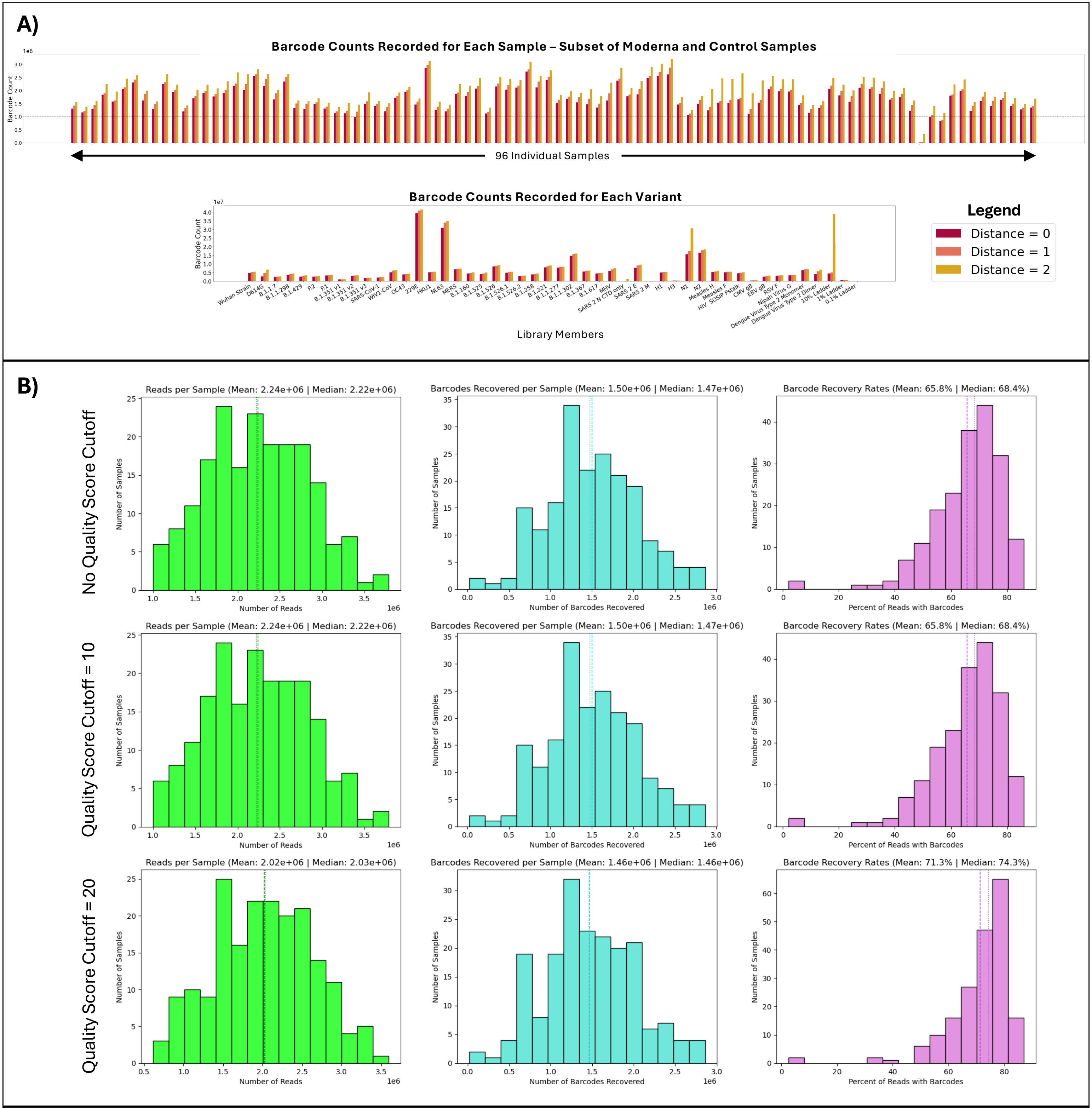


**Supplementary Figure 13.** Effects of altering sequence analysis parameters on ARCADE barcode recovery. **A)** Observed disproportionate increases in barcode counts across samples and variants when allowing mismatches between read barcodes and library barcodes. Barcode counts for a subset of Moderna-vaccinated donor samples are shown (top). Barcode counts for each library member for all Moderna-vaccinated donors are also shown (bottom). The dotted line indicates a count of $10^6$ barcodes. Increasing the Hamming distance between library barcodes and read barcodes from 0 to 1 or 2 increases barcode counts for only certain samples and variants. Some barcodes in the library were only a Hamming distance of 2 away from one another; thus, sequencing errors could allow read barcodes with one or two errors to be assigned incorrectly. A Hamming distance of 0 (indicating an exact barcode match) was chosen for future downstream analyses. **B)** Improved barcode recovery rates from an increased read quality score cutoff. The number of reads per sample, the number of barcodes per sample, and the barcode recovery rate per sample are shown for three quality score cutoffs. Barcode recovery = (barcodes per sample) / (reads per sample). The mean (dashed line) and median (dotted line) are plotted for each graph. The Phred scores for each base in a read were averaged, and reads with average scores under the cutoff were excluded from downstream analyses. A subset of Moderna-vaccinated donor samples are represented by the plots.
